# DecoyOpt: A two-stage pipeline for decoy-aware enzyme discrimination and EC number prediction

**DOI:** 10.64898/2026.11.08.762986

**Authors:** Tiantao Liu, Junwen Deng, Xiaodan Wang, Kepeng Chen, Dutao Zhang, Shirley W. I. Siu

## Abstract

Accurate and efficient EC number prediction is essential for elucidating enzyme function. Although similarity-based and deep learning-based approaches have advanced this task, they often fail to distinguish active enzymes from sequence-similar decoys inactivated by functional-site mutations, resulting in high false-positive rates (FPRs). We propose DecoyOpt, a two-stage pipeline for decoy-aware enzyme discrimination and fourth-level EC number prediction. Specifically, putatively inactive functional-site decoys were generated using ESM3, and 16 protein language models were evaluated across three data splits. In the first stage, a multilayer perceptron trained with classification and ranking losses discriminates enzymes from unannotated proteins and decoys. Then, predicted enzymes are assigned EC numbers using a contrastive learning model. On the most stringent GraphPart 40% split, DecoyOpt achieves an MCC of 0.899 for enzyme discrimination and a D_0_ decoy FPR of 6.17%, outperforming retrained baseline methods. Across all splits, DecoyOpt consistently surpasses these baselines in EC number prediction on the full test set while maintaining performance comparable to CLEAN on the enzyme-only subset. Furthermore, it correctly annotates all four wild-type enzymes and rejects all eight deleterious variants. These results suggest that DecoyOpt provides robust annotation for uncharacterized proteins and functional variants, with potential applications in genomics, synthetic biology, and biocatalysis.

## 1. Introduction

Enzymes catalyze the biochemical reactions that are essential for life. Characterizing their functions is fundamental to understanding cellular metabolism and discovering new biocatalysts. However, genomic and metagenomic sequencing generates protein sequences faster than they can be experimentally characterized, leaving many proteins without reliable functional annotations.^1^ The widely used Enzyme Commission (EC) system classifies enzymes in a four-level hierarchy according to the reactions they catalyze.^2^ Accurate and efficient EC number prediction is critical for elucidating the catalytic roles of enzymes, establishing gene-protein-reaction associations, and reconstructing metabolic networks.^3^

Methods for EC number prediction can be broadly divided into similarity-based and deep learning-based approaches.^3–9^ Similarity-based methods, such as BLASTp, transfer functional annotations from characterized homologs to query sequences.^4^ However, these methods are less reliable for remote homologs. For example, some remote homologs share less than 30% sequence identity while retaining similar functions.^10^ Deep learning-based approaches can be further categorized as classification-based or contrastive learning-based according to their training objectives. The former directly learn mappings from protein features to EC numbers using hierarchical or multi-label classification.^3, 5–8^ For example, DeepECtransformer combines a transformer-based neural network with homology searches, covering 5,360 EC numbers.^5^ However, class imbalance in the training data can limit the ability of these multi-label classification models to identify underrepresented EC numbers. To address this issue, Yu et al. developed CLEAN,^9^ which uses contrastive learning to bring enzyme embeddings with the same EC number closer together and separate embeddings of enzymes with different EC numbers. CLEAN achieves good predictive performance even when distinguishing among thousands of fourth-level EC numbers.

However, CLEAN lacks explicit enzyme/non-enzyme discrimination and assigns EC numbers even to non-enzyme proteins. Models such as DeepEC partly address this limitation by incorporating negative samples during training.^3^ Nevertheless, distinguishing enzymes from negative samples does not necessarily ensure recognition of closely related decoys inactivated by local functional-site mutations. Sartori et al. generated putatively inactive enzyme decoys by systematically mutating functional sites and their neighborhoods.^11^ DeepEC still misclassified many of these decoys as enzymes, resulting in high false-positive rates (FPRs).

In this paper, we propose DecoyOpt, a two-stage pipeline for decoy-aware enzyme discrimination and fourth-level EC number prediction. We first curated EC-annotated enzymes from Swiss-Prot as positive samples and proteins lacking EC numbers (i.e., unannotated proteins) as negative samples.^12^ To generate putatively inactive enzyme decoys, we masked functional sites and used ESM3 to inpaint each masked position with its lowest-probability alternative amino acid (Figure 1a).^13^ To reduce homology bias, we split the data into five folds using GraphPart across three sequence identity thresholds.^14^ Sequence embeddings were first extracted using 16 protein language models (PLMs; Figure 1b) and subsequently used as input to the two-stage pipeline (Figure 1c). In the first stage, a multilayer perceptron (MLP) trained with classification and ranking losses distinguishes enzymes from unannotated proteins and decoys. In the second stage, a model trained with contrastive learning assigns EC numbers to predicted enzymes.

**Figure 1.**
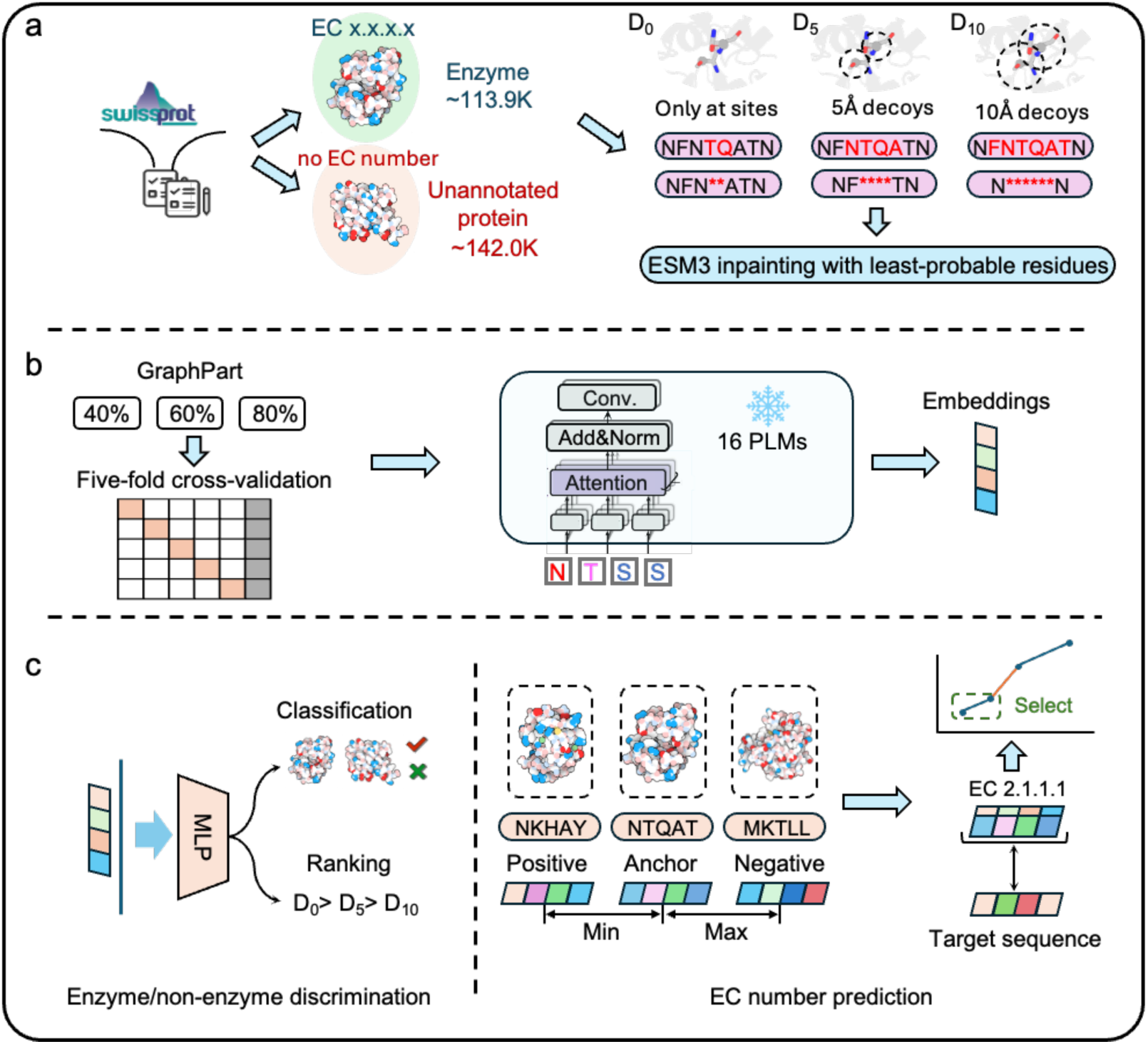
Overview of DecoyOpt. (a) Dataset curation and decoy generation. (b) Dataset split and PLM embedding extraction. (c) Two-stage enzyme/non-enzyme discrimination and fourth-level EC number prediction.

On the most stringent GraphPart 40% split, DecoyOpt achieves an MCC of 0.899 for enzyme/non-enzyme discrimination and a D_0_ decoy FPR of 6.17%, outperforming the retrained DeepEC, ProteInfer, and ECRECer.^3, 6, 7^ Across all three splits, it outperforms these baselines in fourth-level EC number prediction on the full test set while maintaining performance close to CLEAN on the enzyme-only subset, indicating only a small performance loss from the discrimination step. It also correctly assigns EC numbers to all four wild-type enzymes and rejects all eight deleterious variants in the case studies. These results support DecoyOpt’s effectiveness in enzyme/non-enzyme discrimination and EC number annotation, especially in identifying variants impaired by functional-site mutations. This capability broadens the scope of EC number prediction and helps identify deleterious mutations in enzymes, with applications in genomics, synthetic biology, and biocatalysis.

## 2. Methods

### 2.1 Data curation

Enzymes and unannotated proteins were retrieved from Swiss-Prot (release 2026-03).^12^ Enzymes were required to have complete four-level EC numbers, whereas unannotated proteins had no EC number assigned at any level. Both groups were restricted to sequences of 100-1,000 amino acids. Sequence redundancy was reduced using CD-HIT at a 90% sequence identity threshold.^15^ Functional sites of enzymes were obtained from UniProt, including “binding sites”, “active sites”, and “sites”, or predicted using EasIFA when annotations were unavailable.^16^ In total, the final dataset comprised 113,868 enzymes and 141,976 unannotated proteins.

For each enzyme, we generated three putatively inactive decoys. Enzyme structures were retrieved from the AlphaFold-DB (AFDB) or predicted using ESMFold2 when suitable AFDB structures were unavailable.^17, 18^ These structures were used to define three masking regions: functional-site residues alone (D_0_), and those extended to include residues within 5Å (D_5_) or 10Å (D_10_) of any functional-site residue. Following previous work,^11^ we masked each region and used ESM3 (esm3-sm-open-v1) to inpaint it with lowest-probability alternative amino acids. This procedure yielded three corresponding decoys (one each for D_0_, D_5_, and D_10_) for every enzyme in the dataset.

Enzymes and unannotated proteins were partitioned using GraphPart to limit sequence identity between training, validation and test sets.^14^ Each decoy was assigned to the same set as its parent enzyme. Specifically, we used sequence identity thresholds of 40%, 60% and 80%. At each threshold, 10% of sequences were reserved as a fixed test set. The remaining sequences were divided into five folds, with four used for training and one for validation in each fold. The resulting numbers of sequences for each data split are summarized in Table S1.

### 2.2 PLM embeddings

We evaluated 16 pretrained PLMs to select the best one for DecoyOpt. These included ESM-1b,^19^ ESM-1v,^20^ six ESM2 variants (8M, 35M, 150M, 650M, 3B and 15B),^21^ and two ESMC variants (300M and 600M).^22^ We also included five ProtTrans models (ProtBert, ProtBert-BFD, ProtAlBert, ProtT5-XL-BFD and ProtT5-XL-UniRef50)^23^ and the enzyme-specific model EnzGFM.^24^ For each protein sequence, an embedding was obtained from each PLM by mean pooling the outputs of its final hidden layer. The model sizes and embedding dimensions are listed in Table S2.

PLM embeddings were independently projected into two dimensions using UMAP.^25^ UMAP used cosine distance, 100 neighbors, a minimum distance of 0.3 and 500 optimization epochs. The mean silhouette score was calculated to quantify group separation in UMAP space:

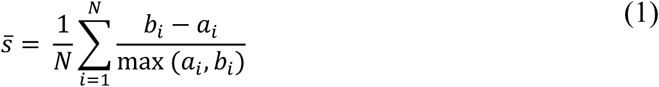

Here, *N* is the number of proteins. In UMAP space, *a_i_* is the mean Euclidean distance to other proteins in the same group, and *b_i_* is that to proteins in the other group.

### 2.3 Model architecture

DecoyOpt comprises two stages that take the same PLM embeddings as input. The first stage uses an MLP to distinguish enzymes from unannotated proteins and decoys. The MLP contains three hidden layers with 512, 512 and 128 units, respectively, followed by a scalar output. Layer normalization is applied to the first two hidden layers, and all three use ReLU activations. Training combines weighted binary cross-entropy (BCE) with a ranking loss:

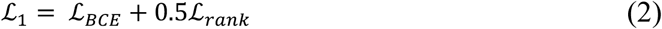

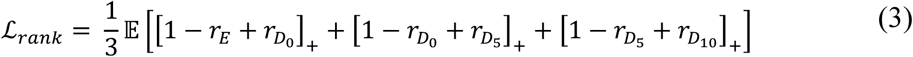

where *r* denotes the output logit, *E* denotes the parent enzyme, and [*x*]_+_ = max (0, *x*). A sigmoid function converts logits into enzyme probabilities.

The second stage follows CLEAN.^9^ A feedforward network maps PLM embeddings to function-aware representations with 128 dimensions. Training uses a triplet margin loss:

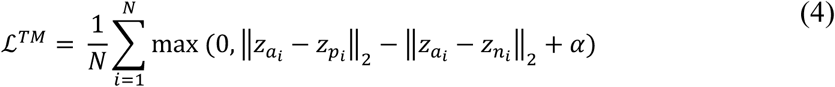

where *N* denotes the number of triplets, *z_x_* denotes the 128-dimensional projected representation of sequence *x*, and *a_i_*, *p_i_* and *n_i_* denote the anchor, positive, and negative, respectively. The margin *α* was set to 1. Positives share an EC number with the anchor, whereas negatives have different EC numbers. Each EC number center is computed by averaging the 128-dimensional projected representations of training enzymes assigned to that class. At inference, query-to-center Euclidean distances are calculated, and fourth-level EC numbers are selected using the maximum-separation method.

In its standard mode, DecoyOpt first distinguishes enzymes from unannotated proteins and decoys. Only sequences classified as enzymes receive EC numbers from the second-stage model. The main hyperparameters of the first- and second-stage models in DecoyOpt are listed in Tables S3 and S4, respectively.

### 2.4 Baseline models

Five baseline models were selected for comparison, including DeepEC, DeepECtransformer, ProteInfer, ECRECer, and EnzBert.^3, 5–8^ All models support enzyme/non-enzyme discrimination and fourth-level EC number prediction.

We first evaluated the original model checkpoints on unannotated proteins and D_0_, D_5_ and D_10_ decoys in our dataset. Given training time and script availability, we further retrained DeepEC, ProteInfer, and ECRECer on our dataset across three identity splits. The training datasets included unannotated proteins and decoys as negative samples. During retraining, the original model architectures and loss functions were retained.

These experiments assessed the ability of the original checkpoints to reject decoys, quantified improvements following retraining, and compared DecoyOpt with the retrained baseline models. Details for baseline models are provided in **Supporting Information Part 1**.

### 2.5 Evaluation metrics

To evaluate enzyme/non-enzyme discrimination, we calculated several metrics using scikit-learn,^26^ including the area under the receiver operating characteristic curve (AUROC), average precision (AP), and macro F1-score. To account for data imbalance, we also calculated the overall Matthews correlation coefficient (MCC) and overall confusion entropy (CEN).^27^ In addition, false-positive rates (FPRs) were calculated separately for four groups of negative samples, comprising unannotated proteins and D_0_, D_5_ and D_10_ decoys:

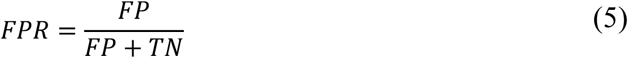

where *TN* and *FP* denote true negatives and false positives, respectively.

Following CLEAN,^9^ we reported weighted precision, weighted recall, and weighted F1-score for fourth-level EC number prediction. We additionally reported macro F1-score and overall MCC. These metrics were calculated on both the full test set and its enzyme-only subset. The latter evaluation assessed the potential loss in EC number prediction performance introduced by first-stage enzyme/non-enzyme discrimination, because enzymes rejected at this stage were counted as false negatives. A summary of the evaluation scenarios and metrics is provided in Table 1.

**Table 1.** Overview of evaluation scenarios and metrics.

| Evaluation scenario | Positive samples (enzymes) | Negative samples | Metrics |
|---|---|---|---|
| Enzyme/non-enzyme discrimination | √ | √ | AUROC, AP, macro F1-score, MCC and CEN |
| FPRs of negative samples | × | √ | FPRs for unannotated proteins, D<sub>0</sub>, D<sub>5</sub> and D<sub>10</sub> |
| EC number prediction on the full test set | √ | √ | Weighted precision, weighted recall, weighted F1-score, macro F1-score, and overall MCC |
| EC number prediction on the enzyme-only subset | √ | × | Same as above |

## 3. Results

### Dataset analysis

We characterized the composition and sequence properties of the dataset (Figure 2). Unannotated proteins are shorter than enzymes, with median lengths of 267 and 350 amino acids, respectively (Figure 2a), whereas their overall amino acid compositions are broadly similar (Figure 2b). The dataset comprises 113,868 enzymes, 141,976 unannotated proteins and one decoy of each type (D_0_, D_5_ and D_10_) per enzyme (Figure 2c). Figure 2d-f show the distributions of maximum sequence identity between each test sequence and the training set at three sequence identity thresholds. All identities remain below their respective thresholds.

**Figure 2.**
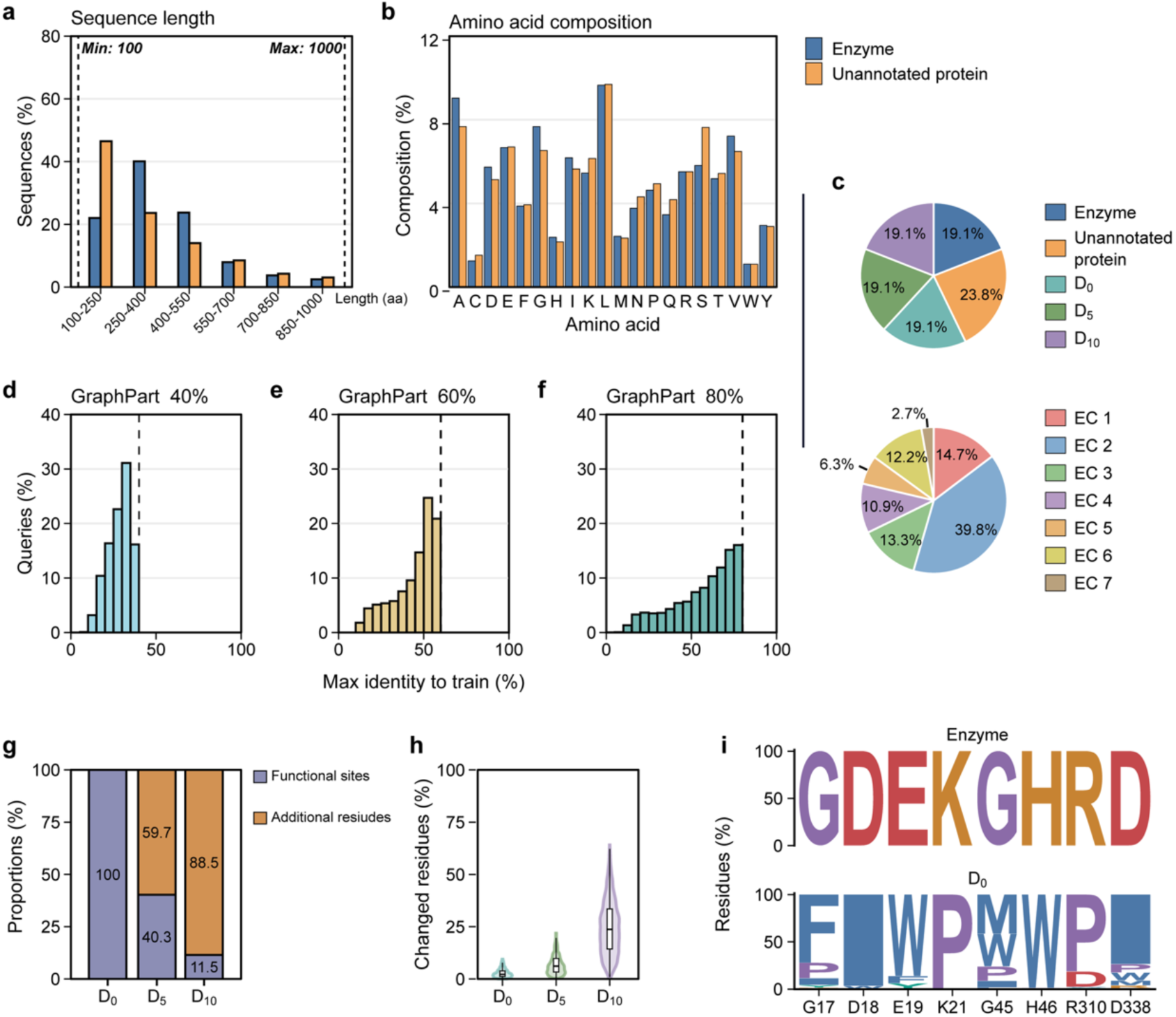
Composition and sequence properties of our dataset. (a) Sequence-length distributions of enzymes and unannotated proteins. (b) Amino acid compositions of enzymes and unannotated proteins. (c) Dataset composition and first-level EC number distribution. (d-f) Training-test sequence identities at identity thresholds of 40%, 60% and 80%. (g) Proportions of functional-site and neighboring residues among the mutated residues in D_0_, D_5_ and D_10_ decoys. (h) Proportions of mutated residues per sequence in D_0_, D_5_ and D_10_ decoys. (i) Conserved functional sites in adenylosuccinate synthase (EC 6.3.4.4; upper panel) and corresponding D_0_ substitutions (lower panel).

Finally, we analyzed the generated decoys (Figure 2g-i). Figure 2g shows the proportions of functional-site residues and additional neighboring residues among the mutated residues in D_0_, D_5_ and D_10_ decoys. In D_10_, neighboring residues substantially outnumber functional-site residues. Figure 2h shows the proportion of mutated residues in each sequence. For D_10_, this proportion averages approximately 25%. Figure 2i shows conserved sites in adenylosuccinate synthase (EC 6.3.4.4; upper panel) and the corresponding substitutions in D0 decoys generated from all enzymes annotated with this EC number (lower panel). Across the D0 decoys, most sites show fewer than three distinct amino acid substitutions.

### PLM embedding analysis

We generated sequence embeddings for enzymes and unannotated proteins using 16 PLMs to visualize their distributions in low-dimensional space. To compare these distributions, we performed UMAP visualization and calculated mean silhouette scores, with higher scores indicating greater class separation in the projected space.

Figure 3 shows the UMAP projections of enzyme and unannotated protein sequences in the GraphPart 40% test set. The mean silhouette scores range from 0.098 to 0.199. ProtBert-BFD achieves the highest score of 0.199, followed by ESM2-t6-8M and EnzGFM. Despite being the largest model evaluated, ESM2-t48-15B yields the lowest score, suggesting that model size alone does not ensure separation between enzymes and unannotated proteins in embedding space and that task-specific training is needed.

**Figure 3.**
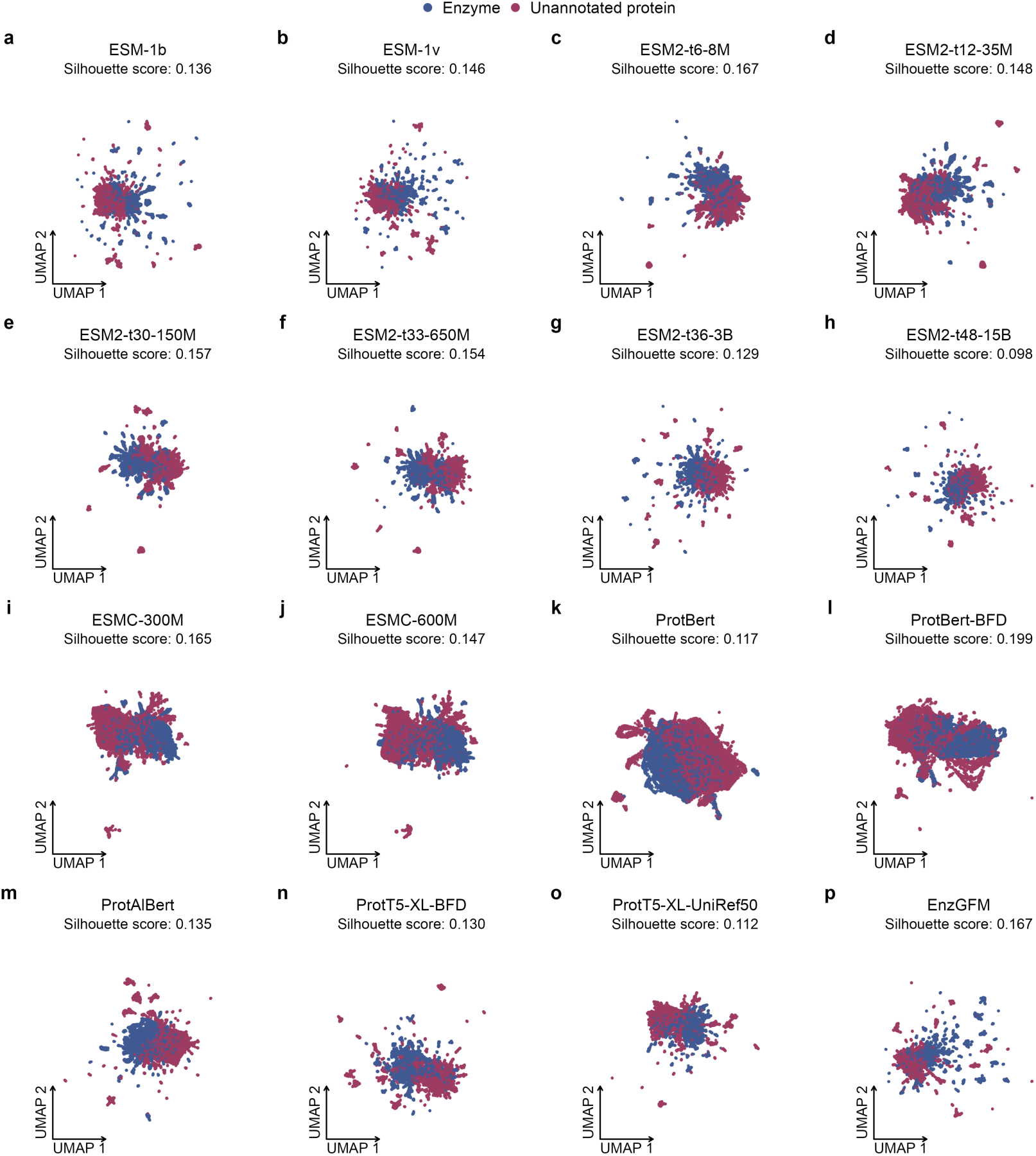
UMAP projections of enzyme and unannotated protein embeddings from 16 PLMs for the GraphPart 40% test set.

### FPRs of original baseline checkpoints

To assess the limitations of existing baselines, we evaluated their publicly released checkpoints on our decoys and unannotated proteins (Figure 4). We selected FPRs as the primary evaluation metric, defined as the proportion of negative sequences falsely predicted as enzymes. All five baselines exhibit FPRs exceeding 75% on D_0_, underscoring their limited ability to distinguish these decoys from enzymes. Under broader mutational perturbations in D_5_ and D_10_, ProteInfer and ECRECer display reduced FPRs, though these remain substantial (>25%). In contrast, FPRs are markedly lower across all baselines on unannotated proteins, with ProteInfer achieving the lowest rate (<2%). Collectively, these findings demonstrate the vulnerability of current models to functional-site mutations, highlighting the necessity of integrating functional-site decoys into both model training and evaluation pipelines.

**Figure 4.**
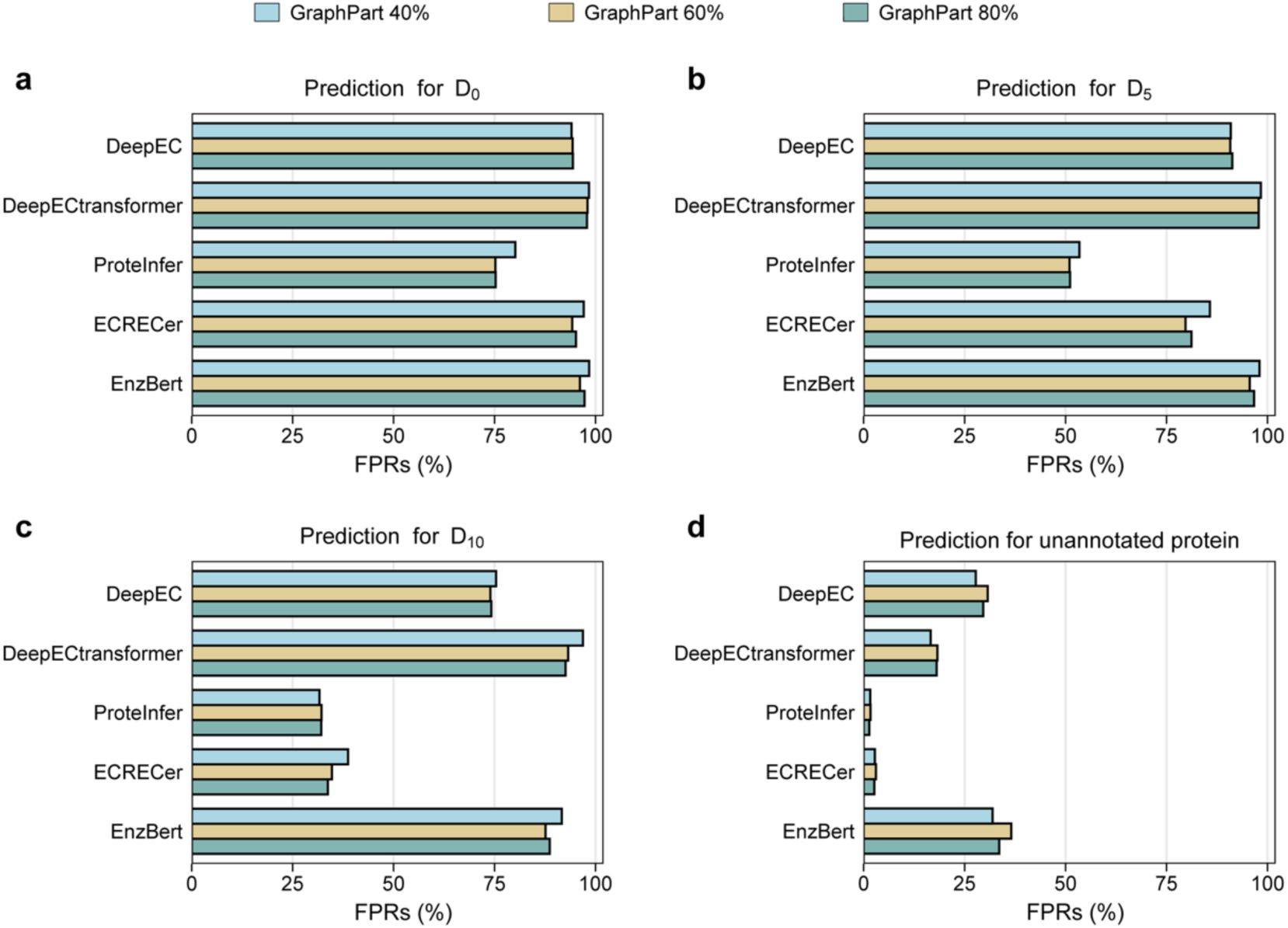
FPRs of five baseline models using their publicly released checkpoints. (a-d) FPRs on D_0_, D_5_ and D_10_ decoys and unannotated proteins, respectively. Colors indicate the test sets from the GraphPart 40%, 60% and 80% splits.

### PLM selection for DecoyOpt

We first evaluated DecoyOpt with embeddings from 16 pretrained PLMs to identify the preferred configuration. For the first-stage enzyme/non-enzyme discrimination, ProtT5-XL-UniRef50 achieves the best performance across all five metrics under all three sequence identity thresholds (Table S5-S7). On the most stringent 40% split, DecoyOpt using ProtT5-XL-UniRef50 achieves a mean MCC of 0.899, outperforming all other PLMs. The largest PLM ESM2-t48-15B and the enzyme-specific PLM EnzGFM achieve mean MCCs of only 0.738 and 0.711, respectively. These results indicate that a larger model size does not necessarily improve predictive performance, consistent with previous findings across other tasks.^28^

We further examined FPRs for decoys and unannotated proteins across the 16 PLMs. As shown in Table S8-S10 and Figure 5, ProtT5-XL-UniRef50, ESMC-600M, and ProtT5-XL-BFD are among the best-performing configurations. On the GraphPart 40% split, ProtT5-XL-BFD and ProtT5-XL-UniRef50 achieve mean FPRs of 6.08% and 6.17% for D_0_ decoys, respectively, both of which are more than 70% lower than the FPR of the original ProteInfer checkpoint. For the more extensively mutated D_10_ decoys, DecoyOpt achieves mean FPRs below 2% with embeddings from each of the 16 PLMs across all three GraphPart splits.

**Figure 5.**
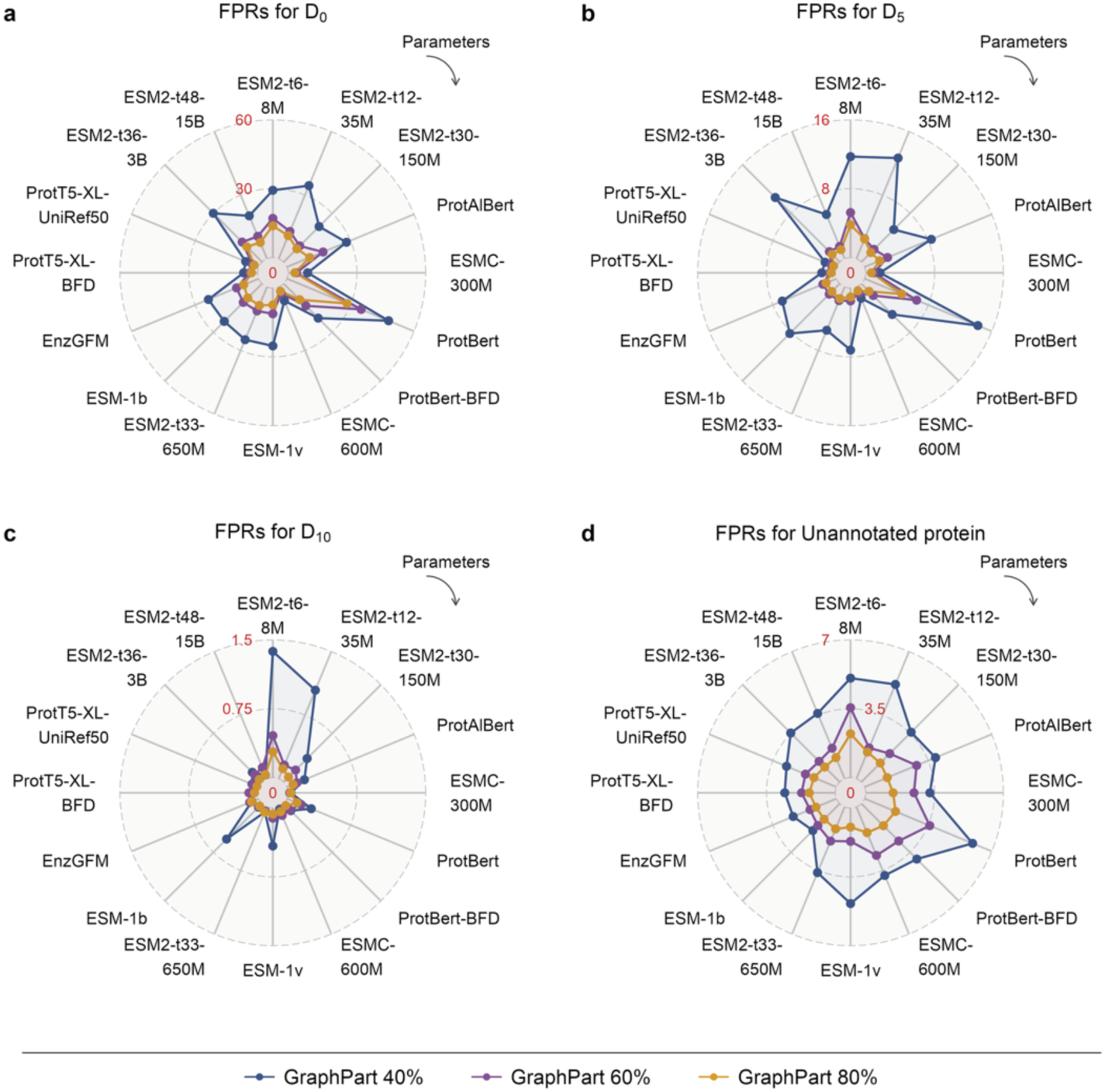
FPRs (%) of DecoyOpt across 16 PLMs and three GraphPart splits. Panels show results for (a) D_0_, (b) D_5_, (c) D_10_, and (d) unannotated proteins. Curves represent five-fold mean FPRs under the GraphPart 40%, 60% and 80% splits. Models are ordered clockwise by increasing model size. Lower values indicate better performance.

We next evaluated DecoyOpt’s fourth-level EC number prediction using each of the 16 PLMs on the full test set and its enzyme-only subset. For evaluation on the full test set, negative samples were treated as a single non-enzyme class and included in metric calculations. ProtT5-XL-UniRef50 shows the strongest overall performance on the full test set (Table S11-S13). It achieves the highest mean weighted precision of 0.905 on the GraphPart 40% split and ranks first across all five metrics on the 60% and 80% splits. This advantage is maintained when the evaluation is restricted to the enzyme-only subset (Table S14-S16). On both the 60% and 80% splits, ProtT5-XL-UniRef50 achieves the highest mean weighted recall, weighted F1-score, macro F1-score, and overall MCC.

Based on its overall performance, we selected ProtT5-XL-UniRef50 as the default PLM for DecoyOpt. Unless otherwise specified, DecoyOpt refers to this configuration.

### Comparison of DecoyOpt with baseline models

We conducted a comprehensive evaluation by comparing DecoyOpt (with ProtT5-XL-UniRef50 embeddings) against retrained versions of DeepEC, ProteInfer, and ECRECer on our dataset. In addition, we retrained a CLEAN model on the enzyme-only subset of our dataset, utilizing triplet margin loss for training and the maximum-separation method for inference.^9^ This baseline provided a reference for assessing the potential loss introduced by first-stage enzyme/non-enzyme discrimination, as true enzymes can be incorrectly rejected as false negatives.

DecoyOpt achieves the best performance in enzyme/non-enzyme discrimination (Figure 6a-c). This advantage is most pronounced under the most challenging GraphPart 40% split, where DecoyOpt achieves a mean MCC of 0.899 and a macro F1-score of 0.949, outperforming the second-best method, ECRECer, by 0.258 and 0.136, respectively. These results demonstrate that DecoyOpt generalizes well to proteins with low sequence identity to the training set. This strong performance extends to FPRs for decoys and unannotated proteins (Figure 6d-f). Specifically, DecoyOpt yields the lowest FPRs across D_0_, D_5_, and D_10_ on the GraphPart 40% split. On the GraphPart 60% and 80% splits, ProteInfer achieves similarly low FPRs for decoys but exhibits substantially higher FPRs for unannotated proteins than DecoyOpt.

**Figure 6.**
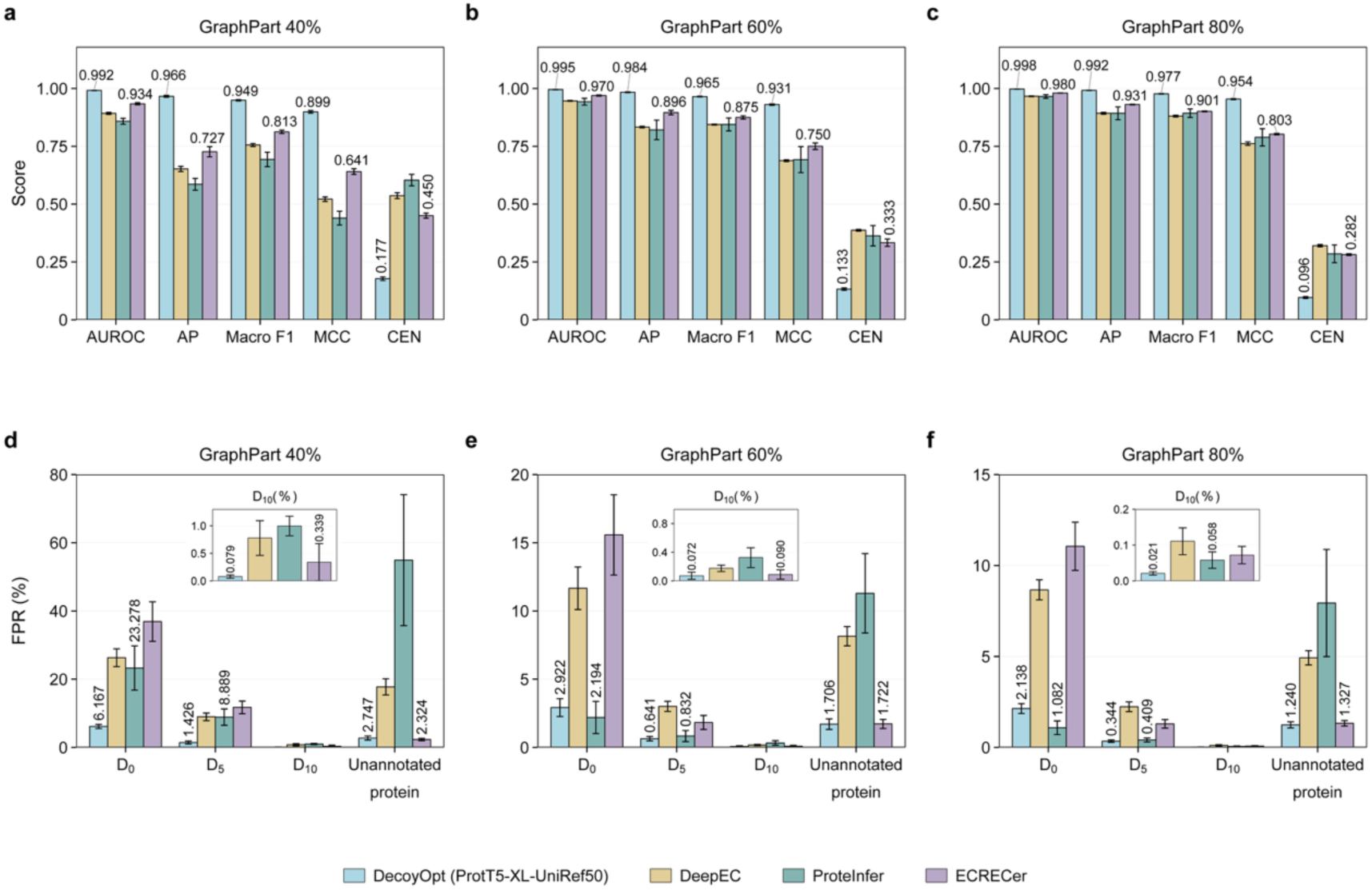
Enzyme/non-enzyme discrimination performance of DecoyOpt and baseline models. DecoyOpt uses ProtT5-XL-UniRef50 embeddings and is compared with retrained DeepEC, ProteInfer and ECRECer. (a-c) AUROC, AP, macro F1-score, MCC and CEN under the GraphPart 40%, 60% and 80% splits, respectively. (d-f) FPRs for D_0_, D_5_, D_10_ and unannotated proteins under the corresponding splits. Bars show five-fold means ± SD. Numerical labels indicate the two best-performing models for each metric.

For EC number prediction on the full test set, DecoyOpt consistently outperforms all three baselines across the five metrics under all three GraphPart splits (Figure 7a-c). On the GraphPart 40% split, it achieves a weighted F1-score of 0.878 and an overall MCC of 0.862, compared with 0.792 for ECRECer and 0.799 for DeepEC, respectively. For EC number prediction on the enzyme-only subset (Figure 7d-f), DecoyOpt continues to outperform DeepEC, ProteInfer and ECRECer, achieving overall MCC values of 0.469, 0.840 and 0.881 under the GraphPart 40%, 60% and 80% splits, respectively. Compared with CLEAN, which predicts EC numbers without enzyme/non-enzyme discrimination, DecoyOpt achieves overall MCC values only 0.038, 0.008 and 0.004 lower under the GraphPart 40%, 60% and 80% splits, respectively. These results show that DecoyOpt enables effective enzyme/non-enzyme discrimination while maintaining EC number prediction performance comparable to that of the state-of-the-art model CLEAN.

**Figure 7.**
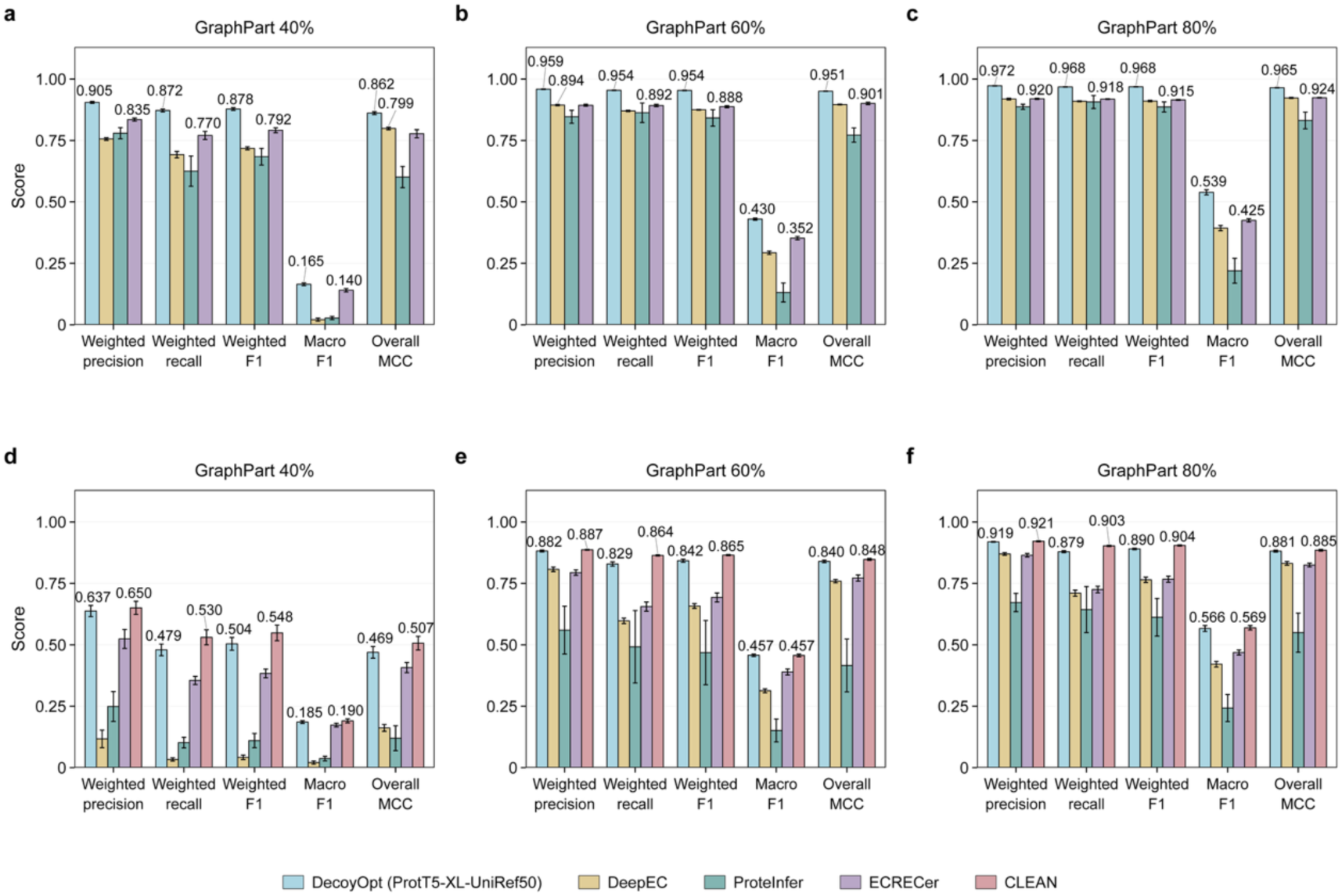
Fourth-level EC number prediction performance on the full test set and enzyme-only subset. (a-c) Weighted precision, weighted recall, weighted F1-score, macro F1-score and overall MCC on the full test set under the GraphPart 40%, 60% and 80% splits, respectively. (d-f) Corresponding results on the enzyme-only subset, including CLEAN. DecoyOpt uses ProtT5-XL-UniRef50 embeddings. Bars show five-fold means ± SD. Numerical labels indicate the two best-performing models for each metric.

### Applications

We consider the principal value of DecoyOpt to lie in distinguishing active enzymes from closely related variants inactivated by local catalytic-site mutations. Accordingly, we selected four enzymes, including IsPETase,^29^ the haloalkane dehalogenases DhlA and DhaA,^30^ and esterase EstD11.^31^ We evaluated four active wild-type enzymes and eight deleterious variants (Figure S1). Despite carrying only one or two amino acid substitutions, these variants were reported to be inactive or nearly inactive and were labeled as non-enzymes in our test set. Table 2 summarizes the predictions of DecoyOpt and the baseline models. In these cases, DecoyOpt recovers the reference EC number for all four wild-type enzymes and correctly classifies all eight deleterious variants as non-enzymes. DeepEC, ProteInfer and ECRECer, retrained on our dataset, show less consistent performance. ECRECer identifies only one of the eight variants as a non-enzyme. Although ProteInfer correctly identifies several deleterious variants, it also misclassifies wild-type enzymes, including DhlA and DhaA, as non-enzymes. These case studies suggest that DecoyOpt can capture the functional effects of deleterious mutations, although systematic validation on larger and more diverse datasets is needed.

**Table 2.** Predictions of DecoyOpt and retrained baseline models for wild-type enzymes and deleterious variants.

| **Entry** | **Label** | **DecoyOpt** | **DeepEC** | **ProteInfer** | **ECRECer** |
|---|---|---|---|---|---|
| IsPETase | 3.1.1.101 | 3.1.1.101 | 3.1.1.101 | 3.1.1.29 | 3.1.1.74 |
| IsPETase-S160A<sup>a</sup> | Non-enzyme | Non-enzyme | 3.1.1.101 | Non-enzyme | 3.1.1.74 |
| IsPETase-D206A | Non-enzyme | Non-enzyme | 3.1.1.101 | 3.1.1.29 | 3.1.1.74 |
| IsPETase-H237A | Non-enzyme | Non-enzyme | 3.1.1.101 | 3.1.1.29 | 3.1.1.74 |
| DhlA | 3.8.1.5 | 3.8.1.5 | 3.8.1.5 | Non-enzyme | 3.8.1.5 |
| DhlA-D124G | Non-enzyme | Non-enzyme | Non-enzyme | Non-enzyme | 3.8.1.5 |
| DhlA-H289Q | Non-enzyme | Non-enzyme | 3.8.1.5 | Non-enzyme | Non-enzyme |
| DhlA-D124A/H289A | Non-enzyme | Non-enzyme | Non-enzyme | Non-enzyme | 3.8.1.5 |
| DhaA | 3.8.1.5 | 3.8.1.5 | 3.8.1.5 | Non-enzyme | 3.8.1.5 |
| DhaA-D106A/H272A | Non-enzyme | Non-enzyme | 3.8.1.5 | Non-enzyme | 3.8.1.5 |
| EstD11 | 3.1.1.1 | 3.1.1.1 | Non-enzyme | 3.1.1.23 | 3.1.1.1 |
| EstD11-S144A | Non-enzyme | Non-enzyme | Non-enzyme | Non-enzyme | 3.1.1.1 |

<sup>*a*</sup>*IsPETase-S160A denotes the inactive IsPETase variant in which serine (S) at position 160 is replaced by alanine (A). The same notation applies to the other variants.*

## 4. Discussion

We propose DecoyOpt, a two-stage pipeline designed to reduce FPRs on negative samples while maintaining high predictive performance in fourth-level EC number prediction. The pipeline incorporates functional-site decoys into enzyme/non-enzyme discrimination and uses contrastive learning to assign EC numbers to predicted enzymes. On the most stringent GraphPart 40% split, DecoyOpt achieves an MCC of 0.899 for enzyme/non-enzyme discrimination and a D_0_ decoy FPR of 6.17%, outperforming all three retrained baselines. For fourth-level EC number prediction across all three splits, DecoyOpt outperforms these baselines on the full test set and maintains performance close to CLEAN on the enzyme-only subset, indicating that the additional enzyme/non-enzyme discrimination step introduces only a small performance loss. DecoyOpt also correctly assigns EC numbers to four wild-type enzymes in the application cases and rejects all eight deleterious variants.

Several limitations remain regarding data availability and the construction of negative samples. Proteins without EC annotations were used as negative samples, although incomplete annotations can leave functional enzymes without assigned EC numbers.^32^ In addition, decoys were generated using ESM3, and their inactivity was inferred from functional-site mutations rather than measured directly. We therefore call for more high-quality, openly accessible datasets containing experimental measurements of enzyme activity to support reliable model training and evaluation.

Nevertheless, incorporating decoys into the negative samples broadens the range of sequences represented in enzyme/non-enzyme discrimination. We believe that DecoyOpt will be a powerful tool for distinguishing enzymes from non-enzymes, particularly inactive variants with localized deleterious mutations, and for assigning accurate EC numbers to predicted enzymes. These capabilities support more reliable functional annotation of uncharacterized proteins and their variants, with applications in genomics, synthetic biology and biocatalysis.

## Data Availability

The source code for DecoyOpt is publicly available at [GitHub](https://github.com/Tiantao2000/DecoyOpt). The DecoyOpt web server is accessible at https://catalymind.com/tools/DecoyOpt/.

## Supporting Information

Additional Information (figures, tables and methodological details) is available in Supporting information. **Part 1**. Details for baseline models; **Part 2**. Supplementary Tables; **Part 3**. Supplementary Figures.

## Acknowledgments

This project was funded by Macao Polytechnic University with grant number RP/FCA-18/2025. The funders had no role in study design, data collection, and interpretation, or the decision to submit the work for publication. This project is part of the thesis work of T. L. with an internal reference number of s/c fca.ca5e.6034.7.

## Author Contributions

T. L. and J. D. contributed to the conceptualization and writing of the manuscript. X. W. and K. C. contributed to revising the manuscript and data curation. D. Z. contributed to data analysis. Shirley W. I. Siu supervised the study and ensured the overall quality of the work.

## Declaration of Interests

The authors declare no competing financial or non-financial interests.

---

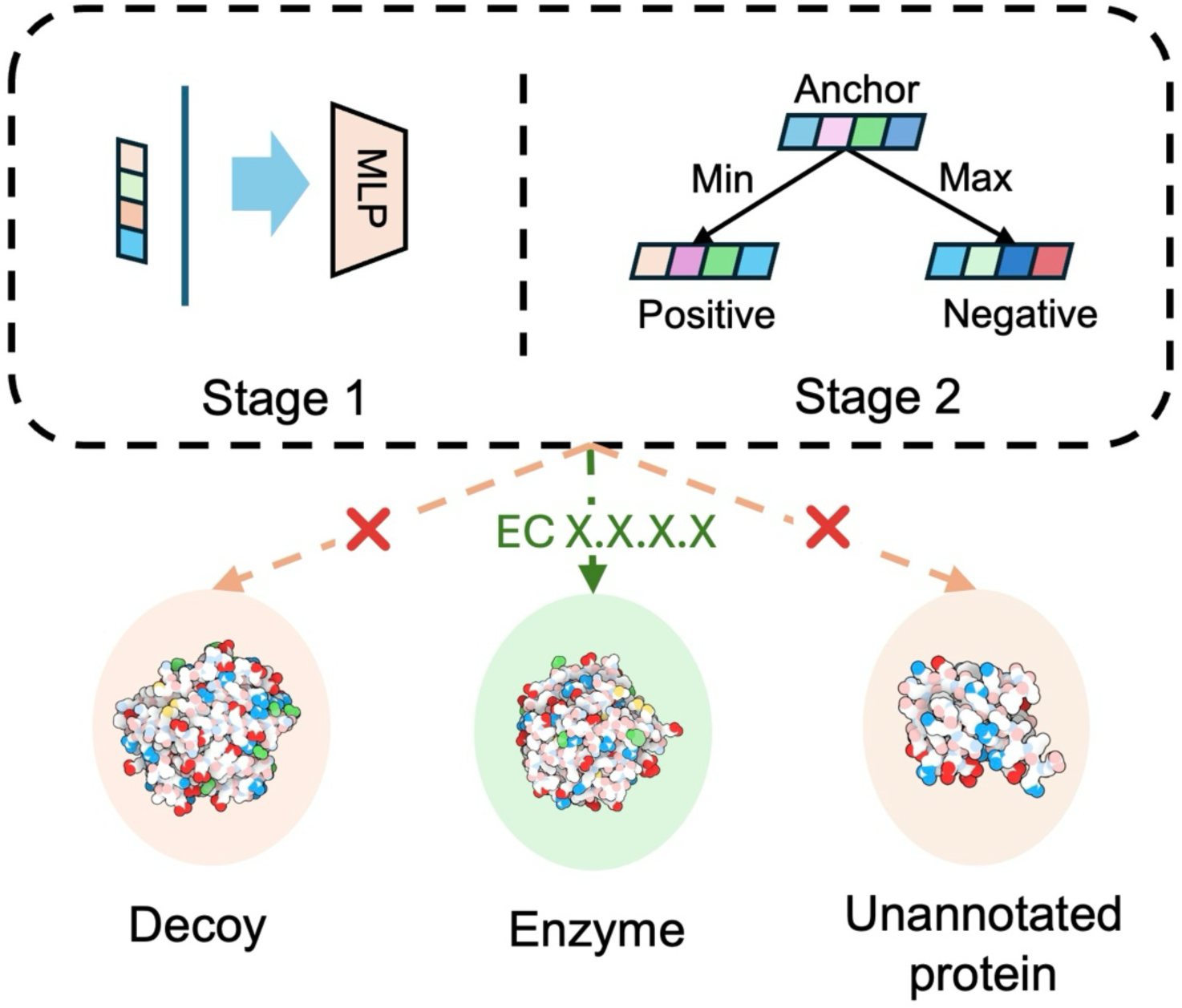
TOC Graphic.

## Notes

### Competing Interest Statement

The authors have declared no competing interest.

## References

(1) Radivojac, P.; Clark, W. T.; Oron, T. R.; Schnoes, A. M.; Wittkop, T.; Sokolov, A.; Graim, K.; Funk, C.; Verspoor, K.; Ben-Hur, A. A large-scale evaluation of computational protein function prediction. Nature methods 2013, 10 (3), 221–227.

(2) McDonald, A. G.; Boyce, S.; Tipton, K. F. ExplorEnz: the primary source of the IUBMB enzyme list. Nucleic acids research 2009, 37 (suppl_1), D593–D597.

(3) Ryu, J. Y.; Kim, H. U.; Lee, S. Y. Deep learning enables high-quality and high-throughput prediction of enzyme commission numbers. Proceedings of the National Academy of Sciences 2019, 116 (28), 13996–14001.

(4) Altschul, S. F.; Gish, W.; Miller, W.; Myers, E. W.; Lipman, D. J. Basic local alignment search tool. Journal of molecular biology 1990, 215 (3), 403–410.

(5) Kim, G. B.; Kim, J. Y.; Lee, J. A.; Norsigian, C. J.; Palsson, B. O.; Lee, S. Y. Functional annotation of enzyme-encoding genes using deep learning with transformer layers. Nature communications 2023, 14 (1), 7370.

(6) Sanderson, T.; Bileschi, M. L.; Belanger, D.; Colwell, L. J. ProteInfer, deep neural networks for protein functional inference. elife 2023, 12, e80942.

(7) Shi, Z.; Deng, R.; Yuan, Q.; Mao, Z.; Wang, R.; Li, H.; Liao, X.; Ma, H. Enzyme commission number prediction and benchmarking with hierarchical dual-core multitask learning framework. Research 2023, 6, 0153.

(8) Buton, N.; Coste, F.; Le Cunff, Y. Predicting enzymatic function of protein sequences with attention. Bioinformatics 2023, 39 (10), btad620.

(9) Yu, T.; Cui, H.; Li, J. C.; Luo, Y.; Jiang, G.; Zhao, H. Enzyme function prediction using contrastive learning. Science 2023, 379 (6639), 1358–1363.

(10) Gerlt, J. A.; Babbitt, P. C. Can sequence determine function? Genome biology 2000, 1 (5), reviews0005. 0001.

(11) Sartori, J.; Guimaraes, A. C. R.; Machado, L. d. A. A novel benchmark dataset for enzyme function prediction reveals the limitations of state-of-the-art models. bioRxiv 2026, 2026.2008. 2021.746242.

(12) UniProt: the universal protein knowledgebase in 2025. Nucleic acids research 2025, 53 (D1), D609–D617.

(13) Hayes, T.; Rao, R.; Akin, H.; Sofroniew, N. J.; Oktay, D.; Lin, Z.; Verkuil, R.; Tran, V. Q.; Deaton, J.; Wiggert, M. Simulating 500 million years of evolution with a language model. Science 2025, 387 (6736), 850–858.

(14) Teufel, F.; Gíslason, M. H.; Almagro Armenteros, J. J.; Johansen, A. R.; Winther, O.; Nielsen, H. GraphPart: homology partitioning for biological sequence analysis. NAR genomics and bioinformatics 2023, 5 (4), lqad088.

(15) Li, W.; Godzik, A. Cd-hit: a fast program for clustering and comparing large sets of protein or nucleotide sequences. Bioinformatics 2006, 22 (13), 1658–1659.

(16) Wang, X.; Yin, X.; Jiang, D.; Zhao, H.; Wu, Z.; Zhang, O.; Wang, J.; Li, Y.; Deng, Y.; Liu, H. Multi-modal deep learning enables efficient and accurate annotation of enzymatic active sites. Nature Communications 2024, 15 (1), 7348.

(17) Varadi, M.; Anyango, S.; Deshpande, M.; Nair, S.; Natassia, C.; Yordanova, G.; Yuan, D.; Stroe, O.; Wood, G.; Laydon, A. AlphaFold Protein Structure Database: massively expanding the structural coverage of protein-sequence space with high-accuracy models. Nucleic acids research 2022, 50 (D1), D439–D444.

(18) Candido, S.; Hayes, T.; Derry, A.; Rao, R.; Lin, Z.; Verkuil, R.; Wu, B. Z.; Lee, J. S.; Bruguera, E. S.; Keval, J. A. Language modeling materializes a world model of protein biology. bioRxiv 2026, 2026.2006. 2003.729735.

(19) Rives, A.; Meier, J.; Sercu, T.; Goyal, S.; Lin, Z.; Liu, J.; Guo, D.; Ott, M.; Zitnick, C. L.; Ma, J. Biological structure and function emerge from scaling unsupervised learning to 250 million protein sequences. Proceedings of the national academy of sciences 2021, 118 (15), e2016239118.

(20) Meier, J.; Rao, R.; Verkuil, R.; Liu, J.; Sercu, T.; Rives, A. Language models enable zero-shot prediction of the effects of mutations on protein function. Advances in neural information processing systems 2021, 34, 29287–29303.

(21) Lin, Z.; Akin, H.; Rao, R.; Hie, B.; Zhu, Z.; Lu, W.; Smetanin, N.; Verkuil, R.; Kabeli, O.; Shmueli, Y. Evolutionary-scale prediction of atomic-level protein structure with a language model. Science 2023, 379 (6637), 1123–1130.

(22) Team, E. ESM Cambrian: Revealing the mysteries of proteins with unsupervised learning. EvolutionaryScale Website, 2024. https://evolutionaryscale.ai/blog/esm-cambrian (accessed).

(23) Elnaggar, A.; Heinzinger, M.; Dallago, C.; Rehawi, G.; Wang, Y.; Jones, L.; Gibbs, T.; Feher, T.; Angerer, C.; Steinegger, M. ProtTrans: toward understanding the language of life through self-supervised learning. IEEE transactions on pattern analysis and machine intelligence 2021, 44 (10), 7112–7127.

(24) Wang, C.; Li, M.; Geng, S.; Li, W.; Zhou, X.; Wang, Y. G.; Yu, Y.; Wang, T.; Shen, Y. An enzyme-specific protein language model for catalytic property prediction. Nature Communications 2026, 17 (1), 8760. DOI: 10.1038/s41467-026-75283-3.

(25) McInnes, L.; Healy, J.; Melville, J. Umap: Uniform manifold approximation and projection for dimension reduction. arXiv preprint arXiv:1802.03426 2018.

(26) Pedregosa, F.; Varoquaux, G.; Gramfort, A.; Michel, V.; Thirion, B.; Grisel, O.; Blondel, M.; Prettenhofer, P.; Weiss, R.; Dubourg, V. Scikit-learn: Machine learning in Python. the Journal of machine Learning research 2011, 12, 2825–2830.

(27) Jurman, G.; Riccadonna, S.; Furlanello, C. A Comparison of MCC and CEN Error Measures in Multi-Class Prediction. PLOS ONE 2012, 7 (8), e41882. DOI: 10.1371/journal.pone.0041882.

(28) Hou, C.; Liu, D.; Zafar, A.; Shen, Y. Understanding language model scaling for protein fitness prediction. Nature Computational Science 2026, 6 (7), 778–788. DOI: 10.1038/s43588-026-01010-z.

(29) Joo, S.; Cho, I. J.; Seo, H.; Son, H. F.; Sagong, H.-Y.; Shin, T. J.; Choi, S. Y.; Lee, S. Y.; Kim, K.-J. Structural insight into molecular mechanism of poly (ethylene terephthalate) degradation. Nature communications 2018, 9 (1), 382.

(30) Fibinger, M. P.; Davids, T.; Böttcher, D.; Bornscheuer, U. T. A selection assay for haloalkane dehalogenase activity based on toxic substrates. Applied microbiology and biotechnology 2015, 99 (21), 8955–8962.

(31) Miguel-Ruano, V.; Rivera, I.; Rajkovic, J.; Knapik, K.; Torrado, A.; Otero, J. M.; Beneventi, E.; Becerra, M.; Sánchez-Costa, M.; Hidalgo, A.;, et al. Biochemical and Structural Characterization of a novel thermophilic esterase EstD11 provide catalytic insights for the HSL family. Computational and Structural Biotechnology Journal 2021, 19, 1214–1232. DOI: 10.1016/j.csbj.2021.01.047.

(32) de Crécy-Lagard, V.; Dias, R.; Sexson, N.; Friedberg, I.; Yuan, Y.; Swairjo, M. A. Limitations of current machine learning models in predicting enzymatic functions for uncharacterized proteins. G3: Genes, Genomes, Genetics 2025, 15 (10), jkaf169.

