## Supporting Information for "DecoyOpt: A two-stage pipeline for decoy-aware enzyme discrimination and EC number prediction"

### Part 1. Details for baseline models

We selected five baseline models, including DeepEC, DeepECtransformer, ProteInfer, ECRECer and EnzBert.[1-5] DeepEC uses three independent CNNs for enzyme/non-enzyme discrimination and EC number prediction. DeepECtransformer uses transformer layers for sequence encoding and combines neural EC number predictions with homology searches. ProteInfer extracts residue-level representations using residual dilated convolutions. These representations are aggregated by mean pooling, followed by sigmoid outputs for EC number prediction. ECRECer combines protein representations with attention-based GRU models for enzyme/non-enzyme discrimination and EC number prediction. EnzBert fine-tunes ProtBert-BFD[6] and uses the final CLS-token representation for EC number prediction. Its classifier treats EC labels as independent classes and includes an explicit non-enzyme class. The code can be found at:

https://bitbucket.org/kaistsystemsbiology/deepec/,

https://github.com/kaistsystemsbiology/DeepProZyme,

https://github.com/google-research/proteinfer,

https://github.com/kingstdio/ECRECer, and

https://gitlab.inria.fr/nbuton/tfpc.

We first evaluated their original checkpoints on unannotated proteins and $D_0$, $D_5$ and $D_{10}$ decoys in our dataset, using FPRs to quantify false-positive predictions. Then, we retrained DeepEC, ProteInfer, and ECRECer on our dataset across three GraphPart splits. Specifically, we used single-model ProteInfer, without model ensembling or integration with BLAST. ECRECer used a fixed configuration combining embeddings from layer 32 of ESM-1b[7] with a bidirectional GRU and an attention mechanism. Training and evaluation were performed on four NVIDIA A30 GPUs. We used the original hyperparameter settings unless otherwise specified.

### Part 2. Supplementary Tables

**Table S1** Dataset composition across GraphPart sequence identity thresholds.

| | 40% | 60% | 80% |
|---|---|---|---|
| Enzymes | 102,495/11,373[a] | 102,480/11,388 | 102,481/11,387 |
| Unannotated proteins | 127,778/14,198 | 127,787/14,189 | 127,778/14,198 |
| Decoys[b] | 307,485/34,119 | 307,440/34,164 | 307,443/34,161 |
| Total | 537,758/59,690 | 537,707/59,741 | 537,702/59,746 |

[a] *Values are reported as training (five-fold cross-validation)/test.*

[b] *Decoys comprise the $D_0$, $D_5$ and $D_{10}$ variants of enzymes in the corresponding training or test set.*

**Table S2** Model sizes and embedding dimensions of the evaluated PLMs

| PLM | Model size | Embedding dimension |
|---|---|---|
| ESM-1b | 650M | 1,280 |
| ESM-1v | 650M | 1,280 |
| ESM2-t6-8M | 8M | 320 |
| ESM2-t12-35M | 35M | 480 |
| ESM2-t30-150M | 150M | 640 |
| ESM2-t33-650M | 650M | 1,280 |
| ESM2-t36-3B | 3B | 2,560 |
| ESM2-t48-15B | 15B | 5,120 |
| ESMC-300M | 300M | 960 |
| ESMC-600M | 600M | 1,152 |
| ProtBert | 420M | 1,024 |
| ProtBert-BFD | 420M | 1,024 |
| ProtAlBert | 224M | 4,096 |

| | | |
|---|---|---|
| ProtT5-XL-BFD | 3B | 1,024 |
| ProtT5-XL-UniRef50 | 3B | 1,024 |
| EnzGFM | 650M | 2,048 |

**Table S3** Hyperparameters of the first-stage model for distinguishing enzymes

| Hyperparameters | Setting |
|---|---|
| Hidden layer dimensions | 512, 512, 128 |
| Output dimension | 1 |
| Activation function | ReLU |
| Normalization | Layer normalization in the first two hidden layers |
| Dropout rate | 0 |
| Batch size | 640 |
| Training steps | 20,000 |
| Optimizer | Adam |
| Learning rate | 5e-4 |
| Adam betas | (0.9, 0.999) |
| Random seed | 1234 |

**Table S4** Hyperparameters of the second-stage EC number prediction model

| Parameter | Setting |
|---|---|
| Hidden layer dimensions | 512, 512 |
| Output embedding dimension | 128 |
| Activation function | ReLU in both hidden layers |
| Normalization | Layer normalization in both hidden layers |
| Dropout rate | 0.1 |
| Loss function | Triplet margin loss |
| Triplet margin | 1 |
| Batch size | Up to 6,000 triplets |
| Training epochs | 2,000 |
| Hard-negative EC neighbors | 30 |
| Hard-negative refresh interval | 100 epochs |
| Optimizer | Adam |
| Optimizer reset interval | 100 epochs |
| Learning rate | 5e-4 |
| Adam betas | (0.9, 0.999) |
| Random seed | 1234 |
| Distance metric | Euclidean distance |
| EC selection method | Maximum separation |

**Table S5** Enzyme/non-enzyme discrimination performance of DecoyOpt with 16 PLMs under the GraphPart 40% split

| PLM | AUROC ↑ | AP ↑ | Macro F1 ↑ | MCC ↑ | CEN ↓ |
|---|---|---|---|---|---|
| ESM-1b | 0.949±0.002 | 0.773±0.029 | 0.845±0.005 | 0.693±0.009 | 0.397±0.009 |
| ESM-1v | 0.950±0.001 | 0.774±0.017 | 0.845±0.004 | 0.697±0.007 | 0.399±0.007 |
| ESM2-t6-8M | 0.923±0.003 | 0.734±0.008 | 0.797±0.004 | 0.604±0.005 | 0.475±0.007 |
| ESM2-t12-35M | 0.932±0.003 | 0.757±0.014 | 0.803±0.006 | 0.623±0.009 | 0.466±0.010 |
| ESM2-t30-150M | 0.960±0.001 | 0.854±0.011 | 0.864±0.005 | 0.732±0.009 | 0.364±0.010 |
| ESM2-t33-650M | 0.965±0.001 | 0.852±0.012 | 0.868±0.007 | 0.743±0.010 | 0.354±0.010 |
| ESM2-t36-3B | 0.946±0.003 | 0.770±0.019 | 0.832±0.006 | 0.676±0.010 | 0.419±0.009 |
| ESM2-t48-15B | 0.960±0.002 | 0.838±0.022 | 0.867±0.006 | 0.738±0.012 | 0.356±0.012 |
| ESMC-300M | 0.986±0.001 | 0.949±0.003 | 0.930±0.003 | 0.860±0.006 | 0.226±0.008 |
| ESMC-600M | 0.989±0.001 | 0.959±0.005 | 0.939±0.002 | 0.878±0.005 | 0.204±0.006 |
| ProtBert | 0.920±0.002 | 0.683±0.015 | 0.784±0.003 | 0.599±0.003 | 0.490±0.003 |
| ProtBert-BFD | 0.965±0.001 | 0.828±0.008 | 0.877±0.004 | 0.759±0.007 | 0.339±0.007 |
| ProtAlBert | 0.946±0.003 | 0.776±0.019 | 0.837±0.008 | 0.683±0.012 | 0.412±0.013 |
| ProtT5-XL-BFD | 0.990±0.001 | 0.964±0.006 | 0.944±0.004 | 0.888±0.007 | 0.191±0.010 |
| ProtT5-XL-UniRef50 | **0.992±0.001** | **0.966±0.004** | **0.949±0.003** | **0.899±0.006** | **0.177±0.007** |
| EnzGFM | 0.951±0.004 | 0.797±0.027 | 0.853±0.005 | 0.711±0.011 | 0.383±0.009 |

**Table S6** Enzyme/non-enzyme discrimination performance of DecoyOpt with 16 PLMs under the GraphPart 60% split

| PLM | AUROC ↑ | AP ↑ | Macro F1 ↑ | MCC ↑ | CEN ↓ |
|---|---|---|---|---|---|
| ESM-1b | 0.983±0.001 | 0.937±0.004 | 0.913±0.003 | 0.826±0.006 | 0.263±0.007 |
| ESM-1v | 0.984±0.001 | 0.941±0.002 | 0.917±0.001 | 0.833±0.001 | 0.255±0.002 |
| ESM2-t6-8M | 0.961±0.001 | 0.870±0.004 | 0.860±0.002 | 0.722±0.005 | 0.368±0.004 |
| ESM2-t12-35M | 0.972±0.001 | 0.905±0.002 | 0.887±0.003 | 0.775±0.006 | 0.311±0.006 |
| ESM2-t30-150M | 0.983±0.001 | 0.937±0.003 | 0.914±0.002 | 0.829±0.004 | 0.260±0.005 |
| ESM2-t33-650M | 0.983±0.000 | 0.937±0.002 | 0.916±0.001 | 0.832±0.003 | 0.257±0.003 |
| ESM2-t36-3B | 0.982±0.001 | 0.933±0.004 | 0.911±0.002 | 0.822±0.004 | 0.268±0.003 |
| ESM2-t48-15B | 0.986±0.001 | 0.945±0.002 | 0.921±0.001 | 0.843±0.003 | 0.245±0.003 |

| | | | | | |
|---|---|---|---|---|---|
| ESMC-300M | 0.993±0.000 | 0.976±0.002 | 0.954±0.002 | 0.908±0.004 | 0.165±0.005 |
| ESMC-600M | 0.994±0.000 | 0.977±0.003 | 0.959±0.002 | 0.918±0.003 | 0.151±0.005 |
| ProtBert | 0.945±0.001 | 0.785±0.007 | 0.826±0.001 | 0.663±0.001 | 0.431±0.001 |
| ProtBert-BFD | 0.980±0.000 | 0.919±0.003 | 0.906±0.001 | 0.813±0.002 | 0.280±0.003 |
| ProtAlBert | 0.973±0.000 | 0.898±0.003 | 0.888±0.003 | 0.778±0.004 | 0.316±0.007 |
| ProtT5-XL-BFD | 0.995±0.000 | 0.982±0.002 | 0.963±0.001 | 0.926±0.002 | 0.139±0.003 |
| ProtT5-XL-UniRef50 | **0.995±0.000** | **0.984±0.002** | **0.965±0.002** | **0.931±0.003** | **0.133±0.005** |
| EnzGFM | 0.984±0.001 | 0.945±0.003 | 0.919±0.003 | 0.839±0.006 | 0.249±0.008 |

**Table S7** Enzyme/non-enzyme discrimination performance of DecoyOpt with 16 PLMs under the GraphPart 80% split

| PLM | AUROC ↑ | AP ↑ | Macro F1 ↑ | MCC ↑ | CEN ↓ |
|---|---|---|---|---|---|
| ESM-1b | 0.991±0.000 | 0.966±0.001 | 0.938±0.001 | 0.875±0.002 | 0.207±0.003 |
| ESM-1v | 0.991±0.000 | 0.968±0.001 | 0.943±0.001 | 0.885±0.002 | 0.194±0.003 |
| ESM2-t6-8M | 0.974±0.001 | 0.905±0.002 | 0.888±0.001 | 0.776±0.003 | 0.315±0.002 |
| ESM2-t12-35M | 0.982±0.001 | 0.935±0.004 | 0.911±0.003 | 0.823±0.006 | 0.266±0.006 |
| ESM2-t30-150M | 0.990±0.000 | 0.963±0.001 | 0.937±0.002 | 0.874±0.004 | 0.208±0.004 |
| ESM2-t33-650M | 0.990±0.000 | 0.962±0.002 | 0.936±0.002 | 0.872±0.003 | 0.211±0.004 |
| ESM2-t36-3B | 0.990±0.000 | 0.961±0.001 | 0.934±0.001 | 0.867±0.002 | 0.217±0.003 |
| ESM2-t48-15B | 0.993±0.000 | 0.971±0.001 | 0.945±0.001 | 0.890±0.002 | 0.188±0.003 |
| ESMC-300M | 0.996±0.000 | 0.987±0.001 | 0.968±0.001 | 0.935±0.003 | 0.126±0.004 |
| ESMC-600M | 0.997±0.000 | 0.989±0.001 | 0.974±0.001 | 0.947±0.002 | 0.107±0.003 |
| ProtBert | 0.962±0.001 | 0.855±0.004 | 0.855±0.001 | 0.716±0.002 | 0.381±0.002 |
| ProtBert-BFD | 0.989±0.000 | 0.955±0.002 | 0.930±0.002 | 0.860±0.003 | 0.226±0.004 |
| ProtAlBert | 0.986±0.000 | 0.949±0.002 | 0.922±0.001 | 0.845±0.003 | 0.243±0.003 |
| ProtT5-XL-BFD | 0.997±0.000 | 0.990±0.000 | 0.973±0.001 | 0.947±0.002 | 0.108±0.003 |
| ProtT5-XL-UniRef50 | **0.998±0.000** | **0.992±0.000** | **0.977±0.001** | **0.954±0.002** | **0.096±0.004** |
| EnzGFM | 0.992±0.001 | 0.970±0.003 | 0.944±0.003 | 0.888±0.006 | 0.191±0.008 |

**Table S8** FPRs (%) of DecoyOpt on negative samples across 16 PLMs under the GraphPart 40% split

| PLM | $D_0$ ↓ | $D_5$ ↓ | $D_{10}$ ↓ | Unannotated proteins ↓ |
|---|---|---|---|---|
| ESM-1b | 23.303±3.811 | 8.211±1.737 | 0.547±0.085 | **1.935±0.278** |
| ESM-1v | 25.228±2.079 | 7.207±1.033 | 0.413±0.090 | 4.858±1.329 |
| ESM2-t6-8M | 29.329±4.076 | 11.737±2.434 | 1.375±0.295 | 5.057±1.084 |
| ESM2-t12-35M | 34.489±4.335 | 12.670±2.128 | 1.045±0.330 | 5.192±1.357 |
| ESM2-t30-150M | 21.878±2.354 | 5.337±0.980 | 0.362±0.084 | 3.584±1.078 |
| ESM2-t33-650M | 24.926±3.234 | 5.455±1.624 | 0.060±0.020 | 3.617±1.403 |
| ESM2-t36-3B | 30.052±3.019 | 10.606±1.517 | 0.149±0.091 | 3.526±0.837 |
| ESM2-t48-15B | 20.234±3.463 | 5.559±1.648 | 0.044±0.046 | 3.596±1.624 |
| ESMC-300M | 8.525±1.315 | 1.644±0.567 | 0.032±0.008 | 3.261±0.557 |
| ESMC-600M | 6.529±0.606 | **1.424±0.361** | **0.026±0.012** | 3.774±0.432 |
| ProtBert | 48.001±2.180 | 14.255±1.326 | 0.287±0.103 | 5.944±0.792 |
| ProtBert-BFD | 21.293±1.354 | 5.072±0.383 | 0.109±0.035 | 3.991±0.928 |
| ProtAlBert | 28.107±3.216 | 8.392±1.259 | 0.206±0.081 | 3.903±1.095 |
| ProtT5-XL-BFD | **6.081±1.309** | 1.569±0.535 | 0.042±0.038 | 2.571±0.766 |
| ProtT5-XL-UniRef50 | 6.167±0.595 | 1.426±0.431 | 0.079±0.030 | 2.747±0.583 |
| EnzGFM | 23.760±2.255 | 6.814±1.636 | 0.081±0.066 | 2.379±0.562 |

**Table S9** FPRs (%) of DecoyOpt on negative samples across 16 PLMs under the GraphPart 60% split

| PLM | $D_0$ ↓ | $D_5$ ↓ | $D_{10}$ ↓ | Unannotated proteins ↓ |
|---|---|---|---|---|
| ESM-1b | 11.582±1.238 | 1.728±0.320 | 0.056±0.039 | 1.574±0.265 |
| ESM-1v | 11.207±0.987 | 1.491±0.083 | 0.111±0.019 | 1.693±0.258 |
| ESM2-t6-8M | 17.067±1.351 | 5.232±0.548 | 0.457±0.099 | 3.553±0.601 |
| ESM2-t12-35M | 12.857±0.821 | 2.511±0.251 | 0.160±0.034 | 1.697±0.320 |
| ESM2-t30-150M | 9.921±0.944 | 2.078±0.433 | 0.184±0.035 | 2.044±0.185 |
| ESM2-t33-650M | 11.335±1.097 | 1.719±0.223 | 0.047±0.015 | 1.880±0.423 |
| ESM2-t36-3B | 12.269±0.594 | 1.705±0.147 | 0.114±0.061 | 1.484±0.214 |
| ESM2-t48-15B | 10.551±0.893 | 1.540±0.302 | 0.132±0.024 | 1.682±0.542 |
| ESMC-300M | 3.600±0.436 | 0.926±0.094 | **0.018±0.006** | 2.450±0.395 |
| ESMC-600M | 2.954±0.235 | 0.803±0.111 | 0.100±0.038 | 2.670±0.249 |
| ProtBert | 35.032±1.920 | 6.544±0.523 | 0.167±0.058 | 3.584±0.446 |
| ProtBert-BFD | 13.702±0.887 | 1.965±0.304 | 0.097±0.039 | 2.691±0.515 |
| ProtAlBert | 17.037±3.128 | 2.882±0.722 | 0.118±0.043 | 2.863±0.312 |
| ProtT5-XL-BFD | 3.391±0.154 | **0.641±0.052** | 0.095±0.044 | 1.714±0.258 |
| ProtT5-XL-UniRef50 | **2.922±0.656** | 0.641±0.163 | 0.072±0.049 | 1.706±0.390 |
| EnzGFM | 10.501±1.817 | 1.702±0.390 | 0.107±0.047 | **1.455±0.147** |

**Table S10** FPRs (%) of DecoyOpt on negative samples across 16 PLMs under the GraphPart 80% split

| PLM | $D_0$ ↓ | $D_5$ ↓ | $D_{10}$ ↓ | Unannotated proteins ↓ |
|---|---|---|---|---|
| ESM-1b | 8.828±0.557 | 1.294±0.214 | 0.037±0.021 | 1.124±0.201 |
| ESM-1v | 7.340±0.642 | 1.024±0.103 | 0.065±0.029 | **0.966±0.102** |
| ESM2-t6-8M | 13.823±0.498 | 3.813±0.311 | 0.281±0.048 | 2.227±0.166 |
| ESM2-t12-35M | 10.856±0.972 | 2.478±0.238 | 0.118±0.039 | 1.464±0.131 |
| ESM2-t30-150M | 8.143±0.288 | 1.554±0.173 | 0.076±0.026 | 1.368±0.144 |
| ESM2-t33-650M | 8.710±0.355 | 1.460±0.198 | 0.063±0.039 | 1.211±0.152 |
| ESM2-t36-3B | 9.363±0.763 | 1.261±0.221 | 0.040±0.013 | 1.080±0.134 |
| ESM2-t48-15B | 7.667±0.572 | 1.133±0.278 | 0.047±0.018 | 1.161±0.123 |
| ESMC-300M | 2.938±0.485 | 0.652±0.072 | 0.028±0.010 | 1.390±0.397 |
| ESMC-600M | **2.064±0.222** | 0.481±0.072 | 0.054±0.023 | 1.414±0.129 |
| ProtBert | 28.397±1.057 | 4.667±0.370 | 0.118±0.053 | 1.700±0.231 |
| ProtBert-BFD | 9.917±0.396 | 1.268±0.184 | 0.030±0.018 | 1.589±0.380 |
| ProtAlBert | 10.719±0.284 | 1.890±0.237 | 0.072±0.031 | 1.242±0.182 |
| ProtT5-XL-BFD | 2.656±0.267 | 0.474±0.101 | 0.033±0.017 | 1.321±0.250 |
| ProtT5-XL-UniRef50 | 2.138±0.271 | **0.344±0.068** | **0.021±0.005** | 1.240±0.167 |
| EnzGFM | 7.103±0.671 | 1.298±0.234 | 0.084±0.022 | 1.127±0.194 |

**Table S11** Fourth-level EC number prediction performance of DecoyOpt on the full test set under the GraphPart 40% split

| PLM | Weighted precision ↑ | Weighted recall ↑ | Weighted F1 ↑ | Macro F1 ↑ | Overall MCC ↑ |
|---|---|---|---|---|---|
| ESM-1b | 0.864±0.004 | 0.825±0.011 | 0.834±0.008 | 0.147±0.009 | 0.812±0.013 |
| ESM-1v | 0.866±0.003 | 0.818±0.006 | 0.834±0.005 | 0.145±0.007 | 0.806±0.007 |
| ESM2-t6-8M | 0.824±0.006 | 0.757±0.012 | 0.779±0.004 | 0.068±0.004 | 0.739±0.014 |
| ESM2-t12-35M | 0.847±0.005 | 0.776±0.013 | 0.799±0.007 | 0.106±0.004 | 0.760±0.015 |
| ESM2-t30-150M | 0.872±0.004 | 0.821±0.005 | 0.835±0.003 | 0.121±0.004 | 0.805±0.007 |
| ESM2-t33-650M | 0.877±0.007 | 0.825±0.009 | 0.841±0.006 | 0.133±0.007 | 0.812±0.011 |
| ESM2-t36-3B | 0.862±0.006 | 0.803±0.009 | 0.819±0.007 | 0.141±0.005 | 0.788±0.010 |
| ESM2-t48-15B | 0.875±0.006 | 0.834±0.008 | 0.844±0.005 | 0.146±0.006 | 0.821±0.011 |
| ESMC-300M | 0.886±0.004 | 0.847±0.005 | 0.855±0.005 | 0.123±0.004 | 0.836±0.005 |
| ESMC-600M | 0.891±0.003 | 0.842±0.003 | 0.851±0.003 | 0.116±0.004 | 0.828±0.004 |
| ProtBert | 0.841±0.005 | 0.737±0.007 | 0.775±0.004 | 0.082±0.003 | 0.718±0.008 |
| ProtBert-BFD | 0.865±0.004 | 0.806±0.006 | 0.824±0.003 | 0.098±0.003 | 0.790±0.006 |
| ProtAlBert | 0.850±0.005 | 0.785±0.012 | 0.807±0.006 | 0.088±0.002 | 0.771±0.013 |

| | | | | | |
|---|---|---|---|---|---|
| ProtT5-XL-BFD | 0.903±0.005 | **0.875±0.009** | **0.880±0.007** | 0.159±0.008 | **0.864±0.009** |
| ProtT5-XL-UniRef50 | **0.905±0.004** | 0.872±0.006 | 0.878±0.005 | **0.165±0.006** | 0.862±0.006 |
| EnzGFM | 0.868±0.003 | 0.831±0.008 | 0.840±0.006 | 0.140±0.005 | 0.821±0.010 |

**Table S12** Fourth-level EC number prediction performance of DecoyOpt on the full test set under the GraphPart 60% split

| PLM | Weighted precision ↑ | Weighted recall ↑ | Weighted F1 ↑ | Macro F1 ↑ | Overall MCC ↑ |
|---|---|---|---|---|---|
| ESM-1b | 0.931±0.002 | 0.924±0.002 | 0.923±0.002 | 0.393±0.003 | 0.920±0.002 |
| ESM-1v | 0.933±0.001 | 0.927±0.001 | 0.927±0.001 | 0.398±0.005 | 0.924±0.001 |
| ESM2-t6-8M | 0.889±0.003 | 0.874±0.002 | 0.875±0.001 | 0.246±0.002 | 0.863±0.003 |
| ESM2-t12-35M | 0.908±0.002 | 0.905±0.002 | 0.901±0.002 | 0.314±0.004 | 0.898±0.003 |
| ESM2-t30-150M | 0.928±0.001 | 0.922±0.002 | 0.921±0.001 | 0.359±0.005 | 0.917±0.002 |
| ESM2-t33-650M | 0.933±0.001 | 0.925±0.001 | 0.925±0.001 | 0.389±0.005 | 0.921±0.001 |
| ESM2-t36-3B | 0.930±0.001 | 0.923±0.001 | 0.923±0.001 | 0.399±0.005 | 0.921±0.001 |
| ESM2-t48-15B | 0.936±0.002 | 0.930±0.001 | 0.929±0.001 | 0.405±0.004 | 0.926±0.002 |
| ESMC-300M | 0.951±0.002 | 0.944±0.001 | 0.945±0.001 | 0.402±0.006 | 0.941±0.001 |
| ESMC-600M | 0.954±0.000 | 0.947±0.001 | 0.947±0.001 | 0.402±0.007 | 0.943±0.001 |
| ProtBert | 0.885±0.002 | 0.854±0.002 | 0.862±0.001 | 0.297±0.004 | 0.847±0.002 |
| ProtBert-BFD | 0.922±0.001 | 0.912±0.002 | 0.912±0.001 | 0.358±0.004 | 0.907±0.003 |
| ProtAlBert | 0.915±0.002 | 0.902±0.005 | 0.904±0.002 | 0.344±0.005 | 0.898±0.005 |
| ProtT5-XL-BFD | 0.957±0.001 | 0.952±0.001 | 0.952±0.001 | 0.410±0.004 | 0.947±0.001 |
| ProtT5-XL-UniRef50 | **0.959±0.001** | **0.954±0.001** | **0.954±0.001** | **0.430±0.004** | **0.951±0.001** |
| EnzGFM | 0.933±0.001 | 0.926±0.002 | 0.925±0.001 | 0.377±0.003 | 0.922±0.003 |

**Table S13** Fourth-level EC number prediction performance of DecoyOpt on the full test set under the GraphPart 80% split

| PLM | Weighted precision ↑ | Weighted recall ↑ | Weighted F1 ↑ | Macro F1 ↑ | Overall MCC ↑ |
|---|---|---|---|---|---|
| ESM-1b | 0.951±0.001 | 0.945±0.001 | 0.945±0.000 | 0.488±0.005 | 0.940±0.001 |
| ESM-1v | 0.954±0.001 | 0.949±0.001 | 0.949±0.001 | 0.505±0.007 | 0.946±0.001 |
| ESM2-t6-8M | 0.911±0.001 | 0.903±0.001 | 0.903±0.001 | 0.350±0.006 | 0.894±0.001 |

| | | | | | |
|---|---|---|---|---|---|
| ESM2-t12-35M | 0.931±0.002 | 0.925±0.001 | 0.924±0.002 | 0.410±0.003 | 0.918±0.002 |
| ESM2-t30-150M | 0.949±0.001 | 0.942±0.001 | 0.943±0.001 | 0.466±0.005 | 0.937±0.001 |
| ESM2-t33-650M | 0.949±0.001 | 0.943±0.001 | 0.944±0.000 | 0.479±0.008 | 0.938±0.001 |
| ESM2-t36-3B | 0.949±0.001 | 0.943±0.001 | 0.943±0.001 | 0.494±0.007 | 0.939±0.001 |
| ESM2-t48-15B | 0.956±0.001 | 0.950±0.001 | 0.951±0.001 | 0.510±0.006 | 0.946±0.001 |
| ESMC-300M | 0.965±0.000 | 0.960±0.001 | 0.960±0.001 | 0.502±0.007 | 0.957±0.001 |
| ESMC-600M | 0.969±0.001 | 0.964±0.001 | 0.964±0.001 | 0.519±0.005 | 0.961±0.001 |
| ProtBert | 0.905±0.001 | 0.885±0.001 | 0.891±0.001 | 0.401±0.006 | 0.881±0.002 |
| ProtBert-BFD | 0.943±0.001 | 0.936±0.002 | 0.937±0.001 | 0.465±0.008 | 0.932±0.002 |
| ProtAlBert | 0.941±0.001 | 0.933±0.001 | 0.935±0.001 | 0.457±0.008 | 0.930±0.001 |
| ProtT5-XL-BFD | 0.970±0.000 | 0.965±0.001 | 0.965±0.001 | 0.517±0.005 | 0.961±0.001 |
| ProtT5-XL-UniRef50 | **0.972±0.001** | **0.968±0.001** | **0.968±0.001** | **0.539±0.010** | **0.965±0.001** |
| EnzGFM | 0.954±0.002 | 0.947±0.002 | 0.948±0.002 | 0.471±0.007 | 0.942±0.002 |

**Table S14** Fourth-level EC number prediction performance of DecoyOpt on the enzyme-only subset under the GraphPart 40% split

| PLM | Weighted precision ↑ | Weighted recall ↑ | Weighted F1 ↑ | Macro F1 ↑ | Overall MCC ↑ |
|---|---|---|---|---|---|
| ESM-1b | **0.646±0.018** | 0.462±0.030 | 0.497±0.028 | 0.177±0.011 | 0.490±0.031 |
| ESM-1v | 0.635±0.016 | 0.471±0.022 | 0.507±0.022 | 0.180±0.005 | 0.492±0.024 |
| ESM2-t6-8M | 0.511±0.016 | 0.263±0.018 | 0.300±0.016 | 0.089±0.005 | 0.275±0.015 |
| ESM2-t12-35M | 0.614±0.033 | 0.406±0.021 | 0.442±0.021 | 0.141±0.003 | 0.420±0.018 |
| ESM2-t30-150M | 0.633±0.015 | 0.421±0.020 | 0.457±0.019 | 0.151±0.003 | 0.428±0.018 |
| ESM2-t33-650M | 0.638±0.023 | 0.467±0.036 | 0.502±0.034 | 0.169±0.005 | 0.473±0.034 |
| ESM2-t36-3B | 0.643±0.022 | 0.454±0.018 | 0.487±0.022 | 0.180±0.005 | 0.465±0.013 |
| ESM2-t48-15B | 0.641±0.033 | 0.467±0.020 | 0.505±0.022 | 0.180±0.011 | 0.480±0.016 |
| ESMC-300M | 0.579±0.020 | 0.387±0.022 | 0.418±0.027 | 0.139±0.002 | 0.383±0.021 |
| ESMC-600M | 0.576±0.019 | 0.347±0.011 | 0.381±0.014 | 0.133±0.004 | 0.337±0.013 |
| ProtBert | 0.593±0.026 | 0.355±0.012 | 0.404±0.016 | 0.119±0.003 | 0.359±0.012 |
| ProtBert-BFD | 0.547±0.030 | 0.342±0.014 | 0.379±0.008 | 0.121±0.004 | 0.339±0.013 |
| ProtAlBert | 0.575±0.013 | 0.331±0.007 | 0.379±0.010 | 0.111±0.003 | 0.342±0.012 |
| ProtT5-XL-BFD | 0.635±0.023 | **0.492±0.028** | **0.517±0.033** | 0.177±0.008 | 0.485±0.027 |

| | | | | | |
|---|---|---|---|---|---|
| ProtT5-XL-UniRef50 | 0.637±0.023 | 0.479±0.024 | 0.504±0.026 | **0.185±0.006** | 0.469±0.024 |
| EnzGFM | 0.633±0.013 | 0.482±0.026 | 0.510±0.029 | 0.167±0.007 | **0.509±0.031** |

**Table S15** Fourth-level EC number prediction performance of DecoyOpt on the enzyme-only subset under the GraphPart 60% split

| PLM | Weighted precision ↑ | Weighted recall ↑ | Weighted F1 ↑ | Macro F1 ↑ | Overall MCC ↑ |
|---|---|---|---|---|---|
| ESM-1b | 0.880±0.004 | 0.771±0.010 | 0.803±0.008 | 0.440±0.004 | 0.814±0.005 |
| ESM-1v | 0.882±0.003 | 0.781±0.004 | 0.812±0.002 | 0.445±0.002 | 0.824±0.001 |
| ESM2-t6-8M | 0.775±0.003 | 0.636±0.011 | 0.669±0.008 | 0.298±0.004 | 0.673±0.007 |
| ESM2-t12-35M | 0.837±0.002 | 0.698±0.009 | 0.736±0.007 | 0.358±0.005 | 0.748±0.005 |
| ESM2-t30-150M | 0.863±0.002 | 0.756±0.006 | 0.786±0.003 | 0.404±0.007 | 0.791±0.002 |
| ESM2-t33-650M | 0.878±0.004 | 0.774±0.008 | 0.806±0.005 | 0.437±0.005 | 0.815±0.004 |
| ESM2-t36-3B | 0.880±0.004 | 0.773±0.005 | 0.806±0.004 | 0.444±0.006 | 0.819±0.003 |
| ESM2-t48-15B | **0.887±0.003** | 0.789±0.011 | 0.818±0.009 | 0.450±0.007 | 0.828±0.004 |
| ESMC-300M | 0.868±0.003 | 0.798±0.002 | 0.817±0.002 | 0.436±0.004 | 0.817±0.004 |
| ESMC-600M | 0.865±0.003 | 0.807±0.002 | 0.821±0.002 | 0.436±0.006 | 0.820±0.002 |
| ProtBert | 0.854±0.005 | 0.711±0.016 | 0.756±0.012 | 0.384±0.008 | 0.762±0.011 |
| ProtBert-BFD | 0.847±0.004 | 0.746±0.004 | 0.775±0.002 | 0.404±0.003 | 0.781±0.003 |
| ProtAlBert | 0.860±0.005 | 0.740±0.016 | 0.777±0.012 | 0.397±0.008 | 0.786±0.007 |
| ProtT5-XL-BFD | 0.878±0.003 | 0.824±0.005 | 0.837±0.004 | 0.440±0.005 | 0.829±0.004 |
| ProtT5-XL-UniRef50 | 0.882±0.004 | **0.829±0.009** | **0.842±0.006** | **0.457±0.005** | **0.840±0.005** |
| EnzGFM | 0.872±0.003 | 0.766±0.013 | 0.797±0.009 | 0.414±0.003 | 0.803±0.006 |

**Table S16** Fourth-level EC number prediction performance of DecoyOpt on the enzyme-only subset under the GraphPart 80% split

| PLM | Weighted precision ↑ | Weighted recall ↑ | Weighted F1 ↑ | Macro F1 ↑ | Overall MCC ↑ |
|---|---|---|---|---|---|
| ESM-1b | 0.915±0.003 | 0.833±0.006 | 0.860±0.005 | 0.536±0.009 | 0.857±0.004 |
| ESM-1v | 0.918±0.002 | 0.834±0.002 | 0.863±0.002 | 0.543±0.004 | 0.863±0.001 |
| ESM2-t6-8M | 0.833±0.003 | 0.710±0.007 | 0.746±0.005 | 0.410±0.006 | 0.745±0.004 |
| ESM2-t12-35M | 0.882±0.003 | 0.770±0.012 | 0.805±0.009 | 0.464±0.008 | 0.801±0.007 |

| | | | | | |
|---|---|---|---|---|---|
| ESM2-t30-150M | 0.903±0.002 | 0.820±0.003 | 0.847±0.002 | 0.515±0.007 | 0.842±0.002 |
| ESM2-t33-650M | 0.910±0.003 | 0.826±0.002 | 0.854±0.002 | 0.527±0.008 | 0.850±0.002 |
| ESM2-t36-3B | 0.917±0.004 | 0.827±0.008 | 0.858±0.006 | 0.541±0.007 | 0.856±0.007 |
| ESM2-t48-15B | **0.920±0.004** | 0.845±0.007 | 0.871±0.005 | 0.555±0.008 | 0.868±0.005 |
| ESMC-300M | 0.907±0.002 | 0.853±0.005 | 0.868±0.003 | 0.532±0.008 | 0.862±0.004 |
| ESMC-600M | 0.908±0.001 | 0.862±0.003 | 0.875±0.002 | 0.548±0.006 | 0.868±0.002 |
| ProtBert | 0.896±0.003 | 0.760±0.008 | 0.808±0.006 | 0.492±0.010 | 0.814±0.005 |
| ProtBert-BFD | 0.896±0.003 | 0.803±0.004 | 0.833±0.003 | 0.514±0.009 | 0.832±0.004 |
| ProtAlBert | 0.907±0.003 | 0.802±0.006 | 0.838±0.004 | 0.509±0.006 | 0.840±0.002 |
| ProtT5-XL-BFD | 0.915±0.001 | 0.870±0.003 | 0.882±0.002 | 0.548±0.005 | 0.872±0.002 |
| ProtT5-XL-UniRef50 | 0.919±0.002 | **0.879±0.004** | **0.890±0.003** | **0.566±0.012** | **0.881±0.004** |
| EnzGFM | 0.914±0.002 | 0.827±0.005 | 0.855±0.003 | 0.513±0.006 | 0.846±0.003 |

### Part 3. Supplementary Figures

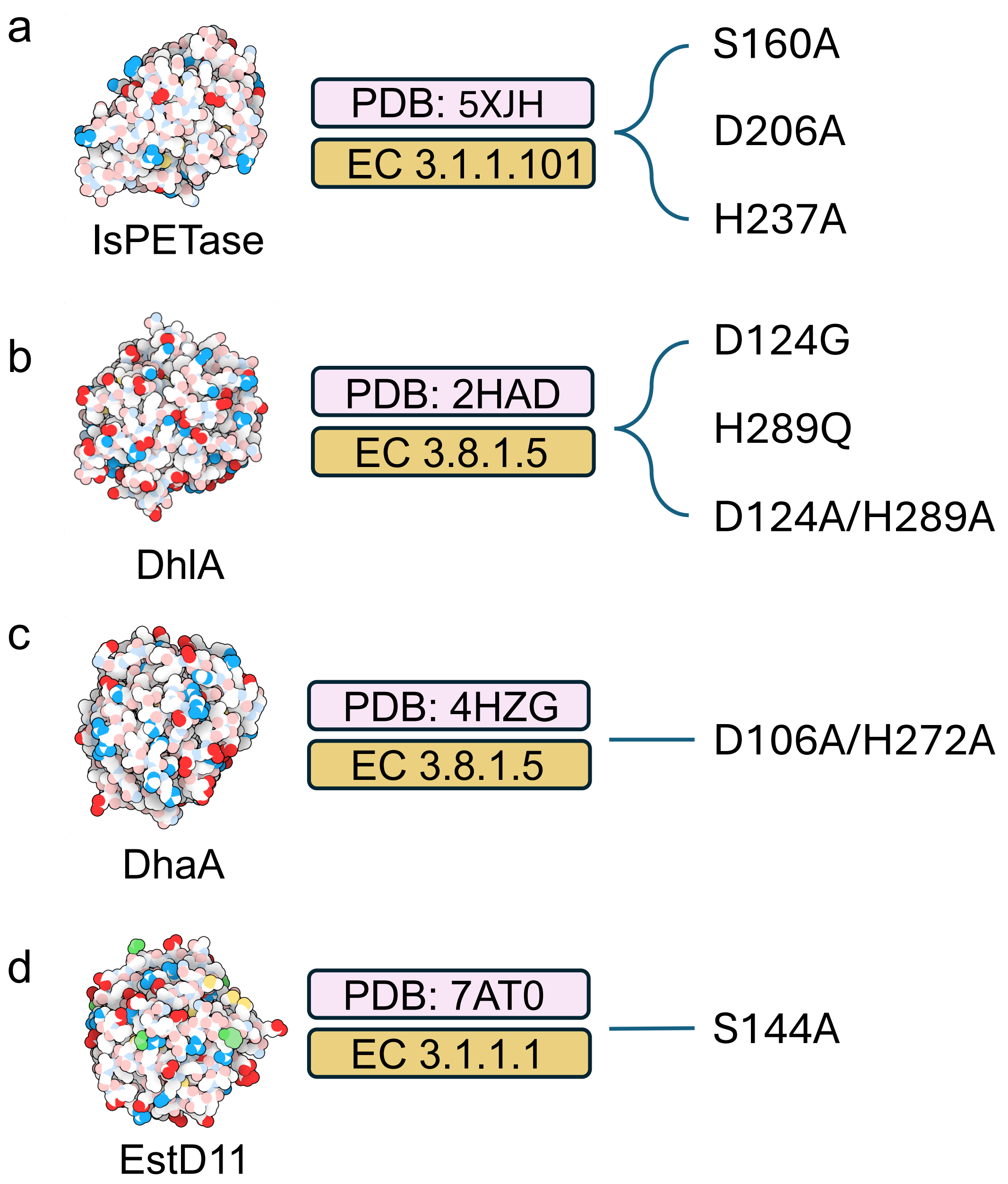


**Figure S1 Wild-type enzymes and their variants.** (a-d) IsPETase (PDB: 5XJH), DhlA (PDB: 2HAD), DhaA (PDB: 4HZG) and EstD11 (PDB: 7AT0), respectively.

### References

(1) Kim, G. B.; Kim, J. Y.; Lee, J. A.; Norsigian, C. J.; Palsson, B. O.; Lee, S. Y. Functional annotation of enzyme-encoding genes using deep learning with transformer layers. *Nature communications* **2023**, *14* (1), 7370.

(2) Ryu, J. Y.; Kim, H. U.; Lee, S. Y. Deep learning enables high-quality and high-throughput prediction of enzyme commission numbers. *Proceedings of the National Academy of Sciences* **2019**, *116* (28), 13996-14001.

(3) Sanderson, T.; Bileschi, M. L.; Belanger, D.; Colwell, L. J. ProteInfer, deep neural networks for protein functional inference. *elife* **2023**, *12*, e80942.

(4) Shi, Z.; Deng, R.; Yuan, Q.; Mao, Z.; Wang, R.; Li, H.; Liao, X.; Ma, H. Enzyme commission number prediction and benchmarking with hierarchical dual-core multitask learning framework. *Research* **2023**, *6*, 0153.

(5) Buton, N.; Coste, F.; Le Cunff, Y. Predicting enzymatic function of protein sequences with attention. *Bioinformatics* **2023**, *39* (10), btad620.

(6) Elnaggar, A.; Heinzinger, M.; Dallago, C.; Rehawi, G.; Wang, Y.; Jones, L.; Gibbs, T.; Feher, T.; Angerer, C.; Steinegger, M. ProtTrans: toward understanding the language of life through self-supervised learning. *IEEE transactions on pattern analysis and machine intelligence* **2021**, *44* (10), 7112-7127.

(7) Rives, A.; Meier, J.; Sercu, T.; Goyal, S.; Lin, Z.; Liu, J.; Guo, D.; Ott, M.; Zitnick, C. L.; Ma, J. Biological structure and function emerge from scaling unsupervised learning to 250 million protein sequences. *Proceedings of the national academy of sciences* **2021**, *118* (15), e2016239118.
